# Promising or problematic? Perceptions of active learning from STEM students with ADHD and specific learning disabilities

**DOI:** 10.1101/2021.12.08.471414

**Authors:** Mariel A. Pfeifer, Julio J. Cordero, Julie Dangremond Stanton

## Abstract

STEM instructors are encouraged to adopt active learning in their courses, yet our understanding of how active learning affects different groups of students is still developing. One group often overlooked in higher education research is students with disabilities. Two of the most commonly occurring disabilities on college campuses are attention-deficit/hyperactivity disorder (ADHD) and specific learning disorders (SLD). We investigated how the incorporation of active-learning practices influences the learning and self-advocacy experiences of students with ADHD and/or SLD (ADHD/SLD) in undergraduate STEM courses. Semi-structured interviews with 25 STEM majors with ADHD/SLD were conducted and data were analyzed using qualitative methods. Most participants perceived themselves to learn best in a STEM course with at least some elements of active learning. Participants described how they perceived active learning to support or hinder their learning and how active learning affected their self-advocacy. Active-learning barriers could be attributed to a combination of instructional factors. These factors included how a particular active-learning practice was implemented within a STEM course and limited awareness of universal design for learning. Defining the supports and barriers perceived by students with ADHD/SLD is a crucial first step in developing more inclusive active-learning STEM courses. Suggestions for research and teaching are provided.

## Introduction

Imagine that you just finished teaching your first STEM class of the term. You are excited. Instead of just lecturing this term, you are ready to try using active learning. Students eager to ask questions approach as you begin to pack your things. You notice one student waiting to speak with you until the rest of the students leave. As the room empties, the student approaches and introduces themselves. They explain that they have been recently diagnosed with ADHD and are planning to use accommodations in your course. They also share that you will soon receive accommodation notifications from the campus disability resource center, which will formally establish these accommodations. You thank the student for sharing this information and the conversation comes to a close. As the student exits the room, you begin to consider, how will the active-learning practices you selected affect students who use accommodations in your course?

In this research article, we begin to address this question. We center the experiences of students with attention-deficit/hyperactivity disorder (ADHD) and specific learning disorders (SLD). We investigate how teaching practices (e.g., lecture versus active learning) within undergraduate STEM courses influence the classroom experiences of students with ADHD and/or SLD. In our study, classroom experiences encompass student perceptions of their own learning and their self-advocacy. Self-advocacy is related to accessing and using accommodations. To build the scholarly context for our study, we present relevant background information about students with disabilities^1^, self-advocacy, and active learning before introducing our guiding frameworks and research questions.

### Students with disabilities

Students qualify to use academic accommodations in a STEM course when they are determined by the campus disability resource center (DRC) to have a disability, as defined by federal laws, such as the Americans with Disabilities Act and Section 504 of the Rehabilitation Act of 1973 and their amendments (Eckes & Ochoa, 2005; Madaus, 2005). Under the Americans with Disabilities Act, disabilities are considered to arise when an individual possesses a condition that has major effects on their daily life. As summarized in our previous work, two of the most reported disabilities on college campuses are attention-deficit/hyperactivity disorder (ADHD) and specific learning disorders (SLD) (Pfeifer et al., 2020, 2021; Raue & Lewis, 2011). ADHD and SLD are considered examples of neurodevelopmental disorders (American Psychological Association, 2013). ADHD has two primary types: primarily inattentive and primarily hyperactive (American Psychological Association, 2013). Features of ADHD in adults can include difficulty maintaining appropriate attention to feelings of restlessness and impulsivity (American Psychological Association, 2013). SLD are diagnosed when an individual shows a marked impairment within a certain academic skill, for example, SLD in reading (dyslexia), SLD in writing (dysgraphia), and SLD in math (dyscalculia) (American Psychological Association, 2013). Features of SLD in reading in adults can include effortful reading and difficulties with reading comprehension. Features of SLD in writing in adults can include difficulty translating thoughts into words. College students with ADHD and SLD share commonalities in the classroom. For example, students with ADHD and SLD can experience similar challenges with motivation, anxiety, and monitoring for understanding (Reaser et al., 2007). Students with ADHD and SLD are frequently studied concurrently because of these similar classroom experiences, ADHD and SLD often co-occur, and because both ADHD and SLD are examples of nonapparent disabilities (DuPaul et al., 2013). Nonapparent disabilities are “impairments with physical and psychological characteristics that are not readily recognized by an onlooker” (Thompson-Ebanks & Jarman, 2018, p. 287). Due to the nonapparent nature of ADHD and SLD, STEM students with ADHD/SLD often need to self-disclose their disability status to others to explain that they qualify for accommodations, or to explain their use of accommodations to those who assume they do not have a disability. This need for self-disclosure is thought to influence the receipt of accommodations. College students with learning disabilities report a lower rate of accommodation receipt compared to students with other types of disabilities (Newman et al., 2011).

### The strengths of STEM students with ADHD/SLD

Most existing research regarding individuals with ADHD and SLD emphasizes the difficulties individuals with these conditions may experience (Antshel, 2018). Few studies examine the strengths of individuals with ADHD and SLD. Hyperfocus, or a “state of heightened, intense focus of any duration, which most likely occurs during activities related to one’s school, hobbies, or screen time” is a potential example strength that STEM students with ADHD possess (Hupfeld et al., 2019). For instance, STEM students with ADHD described that hyperfocusing allowed them to become highly detail oriented when completing course exams (Pfeifer et al., 2020). Participants with ADHD in another study named their high energy levels as an asset for their work (Lasky et al., 2016). This study also investigated how participants with ADHD perceived their work environments to influence symptoms of ADHD. Participants perceived some environments to mitigate their symptoms and they perceived themselves to perform best in jobs that entail “stress or mental challenge, novel or varied tasks, physical labor, hands-on work, or topics of intrinsic interest.” (Lasky et al., 2016, p. 165). These descriptors have the potential to align with the contexts STEM undergraduates encounter, making some undergraduate STEM contexts an environment where students with ADHD may excel. College students with SLD in reading demonstrated enhanced spatial learning compared to students without SLD in reading (Schneps et al., 2012). This finding suggests that students with SLD in reading may be well-suited for STEM pursuits requiring extensive visual processing skills such as radiology, astronomy, and microscopy (Schneps et al., 2012). More research is needed to fully understand the unique strengths of STEM students with ADHD/SLD and how undergraduate STEM courses can be designed to be more compatible with these strengths. Although STEM students with ADHD/SLD likely possess unique strengths that serve them in their STEM majors, many students with ADHD/SLD require academic accommodations to equally access learning opportunities in their courses.

### Self-advocacy

Using accommodations in college involves self-advocacy. Self-advocacy is defined as the “ability to assertively state wants, needs and rights, determine and pursue needed supports” and to obtain and evaluate the needed support with the ultimate goal of conducting affairs independently (Izzo & Lamb, 2002, p. 6; Martin & Marshall, 1995; Pfeifer et al., 2021). Engaging in self-advocacy can be cumbersome for students, yet it is considered vital for student success in college from both students and DRC coordinators (Daly-Cano et al., 2015; Janiga & Costenbader, 2002). Self-advocacy is also the main predictor of GPA for college students with disabilities (e.g., Fleming et al., 2017; Kinney & Eakman, 2017; A. Lombardi et al., 2011). Students with disabilities are less likely to graduate with a STEM degree although they show an equal or sometimes greater interest in STEM majors compared to students without disabilities (Lee, 2020; National Science Foundation, 2019). Thus, understanding self-advocacy and how it is influenced is likely to support the retention of students with disabilities in STEM majors.

A student with self-advocacy is aware of how their disability affects their learning and understands their rights as an individual receiving services under federal law (Test et al., 2005). STEM students with self-advocacy also know the process to obtain accommodations and what accommodations they can request from their college or university, and are cognizant that STEM learning contexts vary, which may influence their accommodation needs (Pfeifer et al., 2020). STEM students with self-advocacy engage in different behaviors, such as communication, leadership, and filling gaps to ensure their success (Pfeifer et al., 2020; Test et al., 2005). Moreover, self-advocacy for STEM students is influenced by beliefs such as view of disability and agency (Pfeifer et al., 2020). A definition of each of these self-advocacy components is presented in Table 1.

**Table 1.** Definitions of self-advocacy components from our model of self-advocacy for students with ADHD/SLD in undergraduate STEM courses. Communication is bolded because it is required for self-advocacy. ^a^ Indicates a definition from Test et al., (2005). ^b^ Indicates a definition from Pfeifer et al., (2020). ADHD is attention-deficit/hyperactivity disorder. SLD is specific learning disorder. DRC is Disability Resource Center.

| **Self-advocacy component** | **Definition** |
| --- | --- |
| Knowledge of self <sup>a</sup> | Awareness of individual strengths and weaknesses as a learner with a disability |
| Knowledge of rights <sup>a</sup> | “Knowing one’s rights as a citizen, as an individual with a disability, and as a student receiving services under federal law” (Test et al., 2005, p. 50) |
| Knowledge of STEM learning context <sup>b</sup> | Awareness that accommodation needs are influenced by the learning environment experienced by students with ADHD/SLD in undergraduate STEM courses |
| Knowledge of accommodations <sup>b</sup> | Awareness of:<br>(1) accommodations that are available to a student with ADHD and/or SLD, and<br>(2) how the accommodation process in college works, including knowledge of the student role, the DRC coordinator role, and the instructor role in the process |
| **Communication <sup>a</sup>** | Communication for the purpose of self-advocacy involves "negotiation, assertiveness, and problem-solving in a variety of situations" (Test et al., 2005, p. 50) |
| Leadership <sup>b</sup> | Taking action for others with diagnosed disabilities to overcome stigma and advocating for peers without formally diagnosed disabilities to be tested to receive academic accommodations |
| Filling gaps <sup>b</sup> | Participants taking action to mitigate a perceived limitation in either their formal accommodations from the DRC, or a perceived limitation in the instructional practices used in a STEM course |
| View of disability <sup>b</sup> | Individual student view of their own disability, and their perceptions of how STEM instructors, and peers view disability and accommodation use in the context of undergraduate STEM courses |
| Agency <sup>b</sup> | An individual belief that an individual student with a disability is responsible for their own accommodations and success in college |

For undergraduate STEM students with ADHD/SLD, self-advocacy is influenced by many internal and external factors (Pfeifer et al., 2021). Internal factors, or aspects within an individual, include the aggregated self-advocacy knowledge an individual holds, their self-advocacy beliefs, and their additional identities, such as racial and gender identities. External factors are aspects beyond the level of an individual student. Example external factors include other individuals such as peers, families, and DRC coordinators, as well as the classroom environment. Classroom environment, which involves the language and actions taken by STEM instructors, is a significant influence upon the self-advocacy of STEM undergraduates with ADHD/SLD. In our previous work, we identified that students perceived STEM instructors to support and, in some cases, hinder self-advocacy (Pfeifer et al., 2021). For instance, STEM instructors could support self-advocacy for students with ADHD/SLD by verbally affirming their use of accommodations in a course. Conversely, STEM instructors could hinder self-advocacy when they lacked knowledge about the instructor’s role in the accommodation process, or through their actions like adopting anti-technology policies in their courses. Given the effect of classroom environment upon self-advocacy, we hypothesized that self-advocacy experiences of students with ADHD/SLD could be influenced by other aspects of the classroom environment. Specifically, we hypothesized that the type of teaching practices used within a STEM course could influence the self-advocacy and learning experiences perceived by students with ADHD/SLD.

### Active learning

Active learning is a term representing many different forms of teaching practices in undergraduate STEM courses (D. Lombardi & Shipley, 2021). Generally, active learning is understood to be the contrast of traditional lecture in which the instructor is talking while students passively listen. Many biology education research studies define active learning by describing the types of active-learning strategies, or practices, present within a certain course (Driessen et al., 2020). For instance, active-learning strategies can include students engaging in clicker questions, group work, worksheets or problem sets, and class discussions. Besides these strategies, other aspects such as “flipped classrooms” and “course structure” are discussed in the context of active learning in undergraduate STEM courses (e.g., Eddy & Hogan, 2014; D. Lombardi & Shipley, 2021). Flipped classrooms, or courses, involve students acquiring knowledge from either pre-recorded lectures or instructor-selected readings or videos outside of class (Bergmann & Sams, 2007; O’Flaherty & Phillips, 2015). In flipped courses, in-class time can be used to address student questions from their outside classwork, or by engaging students in activities and other cognitive work. Course structure can be broadly thought of as the processes and approaches that help students maximize their learning and engagement (Tanner, 2013; Waugh & Andrews, 2020). Some of the benefits of active-learning instruction may result from increased course structure, which can provide students with more practice and feedback and create more inclusive learning environments. For example, Eddy and Hogan (2014) documented that transforming one course from low to moderate structure, specifically the addition of guided-reading questions, preparatory homework, and in-class activities, increased the performance of the entire class while also closing achievement gaps for Black students and first-generation students.

However, not all implementation of active learning leads to increased student learning outcomes. In a population of randomly selected introductory biology instructors, active learning was not associated with enhanced student learning of natural selection (Andrews et al., 2011). The factors contributing to the nuanced outcomes of active learning are not yet understood. Such factors may include issues with implementation of active-learning practices by instructors (e.g., Stains & Vickrey, 2017), as well as the potential for active learning to negatively affect certain groups of students within STEM courses.

Active learning is often assumed to be an inclusive teaching pedagogy with the potential to enhance student learning relative to traditional lecture, especially for higher-order cognitive tasks (e.g., Beichner et al., 2007; Dewsbury & Brame, 2019; Eddy & Hogan, 2014; Freeman et al., 2014; Haak et al., 2011; Theobald et al., 2020). Yet relatively few studies have examined how different student groups are affected by active learning in undergraduate STEM courses. Certain active-learning practices are known to affect students with self-reported anxiety in some classrooms (Brigati et al., 2020; Cooper et al., 2018; Downing et al., 2020; England et al., 2017; Hood et al., 2021). While anxiety can serve as an impetus for students to study, high levels of anxiety can impede academic performance (Downing et al., 2020; Seipp, 1991). Active learning may also affect the experiences of students with LGBTQIA identities in the classroom. Specifically, active learning was found to “increase the relevance of LGBTQIA identities” with the potential to influence feelings of belonging within the classroom environment (Cooper & Brownell, 2016, p. 8). Our understanding of how active learning influences different groups of students is still evolving.

### Active learning and students with disabilities

It is not yet clear how the implementation of active-learning practices within undergraduate STEM courses affects the classroom experiences of students with disabilities who are registered with their campus Disability Resource Center (DRC). Some literature suggests that active learning is challenging for students with disabilities (Gin et al., 2020; Gonzalez, 2017). In an interview study with 37 campus DRC directors, several aspects of active learning were identified that could negatively influence the experiences of STEM undergraduates with disabilities (Gin et al., 2020). For instance, DRC directors stated that students with learning disabilities^2^ could experience difficulty with group work and clicker questions (Gin et al., 2020). We note that DRC directors are likely most familiar with the challenges of active learning, and not necessarily the benefits of active learning for students because they may or may not be meeting with students directly in their day-to-day work. In our own experiences, DRC directors are typically called upon when an accommodation problem needs to be solved, not necessarily when accommodations are working well for students. This study highlights issues about active learning implementation that STEM instructors should know. To complement this work, we need to consider the voices of current STEM students themselves.

Two studies begin to address the need for student voice. One study found students with learning disabilities in undergraduate STEM courses preferred hands-on learning and reported traditional lecture as their least preferred learning method (Cox et al., 2019). The connection between hands-on learning and active learning is not well-defined. But it can be reasonably assumed that some aspects of hands-on learning are similar in nature to active-learning practices in a STEM course. In a separate study of three participants with ADHD in an active-learning physics course, participants reported both benefits and barriers to their learning because of active learning. This study was conducted in a Student-Centered Activities for Large Enrollment Undergraduate Programs (SCALE-UP^3^)-inspired introductory physics course, which used active-learning practices (James et al., 2020). One participant reported that the SCALE-UP course supported their learning because there was “space for being distracted,” or that the nature of the SCALE-UP course allowed them “autonomy in how they learned the material” that was conducive to them as a student with ADHD (James et al., 2020, p. 16). The remaining two participants reported barriers to their learning; the physical arrangement of the classroom was distracting to them, and they were unsure of how to prepare outside of class for the in-class active-learning practices (James et al., 2020). Together, these studies suggest that active learning is likely imparting a more nuanced effect upon students with ADHD and learning disabilities than previously recognized. By more fully understanding this nuance, supports can be optimized and barriers can be minimized, which would likely lead to the generation of more inclusive active-learning STEM courses. Thus, further research is required to more fully understand how implementation of active learning affects students with ADHD and SLD in different institutional contexts and across STEM disciplines.

### Our current study

The purpose of our foundational study is to characterize the learning and self-advocacy experiences of students with ADHD and/or specific learning disorders (SLD) in undergraduate STEM courses that incorporate aspects of active learning, i.e., active-learning practices. Our work is theoretically guided by the social model of disability. In the social model of disability, disability is not due to a biological difference (called an impairment), but rather disability is caused when societal expectations impose difficulty or hardship onto an individual with an impairment (Berghs et al., 2016; Haegele & Hodge, 2016). From the lens of the social model, we predicted that since active-learning practices were likely designed without consideration for the experiences of students with ADHD/SLD, certain active-learning practices could negatively affect our participants. Because of this, participants would likely need to engage in self-advocacy. This study is a component of a larger study regarding the self-advocacy experiences of STEM undergraduates with ADHD and/or SLD (ADHD/SLD) (Pfeifer et al., 2020, 2021). We used the emerging theory from our conceptual model of self-advocacy (*See Table 1 for components*) to define self-advocacy in this work.

The overarching research question (RQ) guiding this study is: How does the implementation of active-learning practices in undergraduate STEM courses affect the perceived learning and self-advocacy experiences of students with ADHD/SLD? We investigated this question by conducting semi-structured interviews with 25 participants. In our study, we address four specific research questions:

RQ1. In what learning environment do students with ADHD/SLD perceive themselves to learn best?
RQ2. What aspects of active learning influence students’ perceptions of learning?
RQ3. How does active learning influence self-advocacy?
RQ4. What recommendations do students with ADHD/SLD offer to instructors to enhance their learning experiences in active-learning STEM courses?

## Methods

Data analyzed for this paper were collected as part of a larger study investigating the self-advocacy experiences of STEM students with ADHD and/or SLD (ADHD/SLD) (Pfeifer et al., 2020, 2021). For this study, we examined a subset of our data that focused on the experiences of our participants in STEM courses with active-learning practices. These data were not included in our previous analyses. We recruited 25 STEM majors with ADHD/SLD during the Fall 2018 and Spring 2019 semesters in partnership with a Disability Resource Center (DRC) in an accessible and confidential manner, as we previously described (Pfeifer et al., 2020, 2021). A summary of participant characteristics is presented in Table 2. All participants provided informed consent and were compensated $20 for completing the study. The study was deemed exempt for IRB review (STUDY00004663). In the following subsections, we describe the context of our study, our positionality, data collection, data analysis, and the trustworthiness of our study

**Table 2.** Summary of participant characteristics. Both indicates both ADHD and specific learning disability (SLD). Race is not reported at the individual level to protect confidentiality. Out of 25 participants, two participants are Black and 23 are white. Year indicates year in college. Fifth and up indicates participants in their fifth year or greater of college.

| **Participant** | **STEM Major** | **Disability** | **Gender** | **Year** |
| --- | --- | --- | --- | --- |
| Allissa | Engineering | ADHD | Woman | Fourth |
| Brett | Engineering | ADHD | Man | Fifth and up |
| Bryce | Life Science | ADHD | Man | Second |
| Carson | Mathematics-Related | Both | Man | Fifth and up |
| Dylan | Life Science | ADHD | Man | Second |
| Elliott | Life Science | ADHD | Man | Fourth |
| Erik | Mathematics-Related | Both | Man | First |
| Felix | Engineering | Both | Man | Fourth |
| Jack | Physical Science or Technology | SLD | Man | First |
| Jessa | Physical Science or Technology | ADHD | Woman | Third |
| Josiah | Life Science | SLD | Man | First |
| Kacey | Life Science | ADHD | Woman | Second |
| Lena | Engineering | ADHD | Woman | Fifth and up |
| Lucas | Life Science | ADHD | Man | Third |
| Mark | Life Science | ADHD | Man | Fifth and up |
| Olen | Life Science | ADHD | Man | Third |
| Penny | Engineering | Both | Woman | Fifth and up |
| Sadie | Engineering | ADHD | Woman | Fifth and up |
| Stella | Physical Science or Technology | Both | Woman | Third |
| Stewart | Life Science | ADHD | Man | Third |
| Therese | Life Science | ADHD | Woman | Third |
| Thomas | Engineering | SLD | Man | Fifth and up |
| Vivian | Life Science | SLD | Woman | Fourth |
| Wren | Life Science | ADHD | Woman | Third |
| Zara | Life Science | SLD | Woman | Third |

### Context of study

Data were collected at a large research-intensive university in the southeastern United States. Prior to data collection for this study, several ongoing university-wide initiatives promoted the adoption of active-learning practices within undergraduate STEM courses. These initiatives included the establishment of a $1 million fund to transform physical classrooms into spaces conducive for active learning (i.e., changing the layout of large lecture halls to promote adoption of group work) as well as the creation of an “intensive summer institute to promote a wider adoption of active-learning pedagogies” (Office of Instruction, 2021). This summer institute was open to all faculty across campus in Summer 2018, prior to data collection. These university-wide initiatives created an appropriate context for our study. That is, many undergraduate STEM courses were using active-learning practices at the time of data collection.

### Positionality of research team

Our research team was comprised of three members. Our relevant experiences are reported in aggregate in an effort to protect confidentiality. At least one or more of our research team was, or were, a STEM major with ADHD/SLD. At least one or more of our research team was, or were, previously a DRC coordinator at a different university than where data collection occurred, and at least one or more of our research team had teaching experience in undergraduate STEM courses.

### Data collection: Screening survey

Participants first completed a brief online screening survey (Supplemental File 1). The purposes of the survey were to: (1) confirm participant eligibility for the study, (2) collect information about participant familiarity with active-learning practices in STEM courses, and (3) customize questions included in the interview protocol (*See Interviews*). The screening survey asked participants to indicate their major, year in school, disability type, and to confirm they were 18 years of age or older. In addition, the survey asked participants to indicate all the STEM courses they were enrolled in or had completed at the time data were collected. A description of active learning was provided for reference in the survey, stating, “Active learning is a type of instruction that instructors in college science, technology, engineering, and mathematics (STEM) courses sometimes use. Active learning may occur when the instructor is not lecturing. Examples of active learning are clicker questions, group work, completing worksheets in class either individually or in a group, class discussions, and student presentations” (*See Supplemental File 1*). Examples of student activities were derived from the student codes of the Classroom Observation Protocol for Undergraduate STEM (COPUS) (Smith et al., 2013). Participants were then asked if their STEM instructor used active-learning practices and to select which practices they remembered their instructor using in their most recent STEM course. Besides the provided example, active-learning practices participants could also select “other” or “don’t remember.” If participants selected “other” they were prompted to describe that practice. After completing the screening survey, participants completed a semi-structured interview typically within 24 to 48 hours.

### Data collection: Interviews

We conducted semi-structured interviews with each participant in person. The average length of the interview was 80 minutes long. Interview questions related to this study are available in Supplemental File 2. The course information collected in the screening survey was used as the basis for Question 1 in the interview protocol. During the interview, 3”x5” notecards were employed as a visual aid to support elicitation efforts. In our preliminary interviews (*See Pfeifer et al., 2020 for a description of these*) we found that the interviewer and the participant often lost track of what specific STEM course we discussed as we moved through a series of interview questions. In our finalized interview protocol, the use of notecards as a visual aid improved the quality of our interview data. Following the interview, participants completed a short demographic survey. Interviews were transcribed by a third-party service, and each resulting transcript was checked for accuracy prior to analysis.

### Data Analysis

We entered into analysis of our transcript data for this study with deep existing knowledge of our data. This familiarity with our data was generated over the course of several years and through two previous in-depth, theory-building analyses of the self-advocacy portions of the interview. Because of our extensive knowledge of our data, structural coding was used to identify relevant segments of the interviews that addressed our overall research question, “How does the implementation of active-learning practices in undergraduate STEM courses affect the perceived learning and self-advocacy experiences of students with ADHD/SLD? Structural coding “both codes and initially categorizes the data corpus to examine comparable segments’ commonalities, differences, and relationships” (Saldaña, 2016, p. 98). Structural coding was performed by one researcher in MaxQDA 2020 and resulted in the identification of relevant transcript segments that were further analyzed.

In the next level of analysis, one researcher reviewed the relevant segments of each interview to generate attribute codes. Attribute codes label vital contextual information about the data and participant characteristics (Saldaña, 2016). The attribute codes in our study were basic demographic information about the participant, along with the name of the STEM course with active-learning practices, and the active-learning practices they named as occurring in this course during the interview. For example, the code “activities reported” was used to summarize the *a priori* active-learning practices that a participant discussed in their interview. We relied on the student activity codes from the COPUS to generate our *a priori* attribute codes of active-learning practices, such as clicker questions, group work, completing worksheets in class either individually or in a group, class discussions, and student presentations (Smith et al., 2013). The STEM course notecards created during the interview were consulted, as needed, during attribute coding. The attribute codes summarized the context needed for subsequent analysis steps by the full research team.

Subsequent analysis involved open or initial coding the relevant interview segments to understand the range of data. One researcher initial coded all 25 interviews, while another researcher initial coded eight interviews. Five of these eight interviews were selected because they reflected the extremes of our data in terms of participant experiences in STEM courses with active-learning practices. The remaining three of eight interviews were selected randomly. After initial coding by both researchers was completed, we met to discuss what questions or hypotheses the data could address in terms of participant perceptions of active-learning practices in their STEM courses. These questions and hypotheses were compiled into a coding matrix (Supplemental File 3). This resulting coding matrix was used by each researcher individually to analyze the interview. After coding a set of four to five interviews, the researchers met to discuss how data were coded and to resolve coding differences. In this meeting a final combined coding matrix was generated that reflected our mutual understanding of the data from a single participant. In this type of analysis, we conducted a mid-level form of holistic coding in combination with in-vivo and descriptive coding to identify themes and sub-themes that emerged from our analysis. One researcher took the lead in compiling the themes and sub-themes into tables and shared the resulting tables with the other researcher. We met to discuss the themes and sub-themes and to resolve any disagreements. Our themes and sub-themes were presented to qualitative researchers who were unfamiliar with our data corpus for feedback and discussion about the resulting themes and sub-themes. This feedback helped us clarify our theme and sub-theme definitions. We also generated many different visual representations of our data to help refine themes and sub-themes. Analytic memos were kept throughout the entire data analysis process to help track and monitor our reactions, decisions, and interpretations.

### Trustworthiness of study

The criteria to assess rigor differs between quantitative and qualitative research. Tracy (2010) provides a model of criteria to guide the assessment of qualitative research. These criteria include worthy-topic, rich rigor, sincerity, credibility, resonance, significant contribution, ethical, and meaningful coherence (Tracy, 2010). In our view, readers are the ultimate judge of some of these criteria, such as worthy-topic, significant contribution, and meaningful coherence. However, we acknowledge that the research team can enhance the transparency (or sincerity) of their work by articulating how their own research endeavored to address the remaining criteria. In terms of rich rigor, our study involved 25 participants, which is a relatively large sample size for a study of its nature. We sought to establish sincerity by engaging in self-reflexivity throughout the study, evident in our analytic memos during study design, data collection, and data analysis. Moreover, we provide transparency in our methods by acknowledging our positionality, intentions, and our challenges in the research process. One example challenge we encountered was keeping track of STEM courses during the preliminary interviews, which we addressed by changing the interview protocol to include notecards as an elicitation aide. We strove for credibility by employing triangulation, including the use of multiple researchers to code the data to consensus. Coding to consensus ensures that all research team viewpoints are considered during data analysis. We consider this a particular strength of our process because our research team included at least one or more researchers who was or were a STEM major with ADHD/SLD. Because our study focused on the experiences of students, we view this as an essential component of our study’s credibility. We further strove for credibility by presenting multiple voices in our results to represent the breadth of our participant’s experiences. In addition to the typical ethical standards of qualitative research, we assigned participants new pseudonyms for this paper that differ from our previous papers to protect confidentiality. Finally, we provide contextual details in our results section. The purpose of these details is to aid readers in finding transferability of our findings to their own contexts where applicable. We address important considerations for transferability to other contexts, in the Limitations section.

### Limitations

We present this section to explain the important factors readers should consider when assessing the transferability of our findings to other contexts. Active learning in our study was purposefully broadly defined in an effort to allow the most salient features of participant-selected active-learning practices to emerge from the interviews. We acknowledge that in our study the “tools, e.g., clickers” and the “actual methodology of active learning” are presented simultaneously (Eddy et al., 2015, p. 2). Yet the accessibility of both active-learning tools and methodologies matter in the overall experiences of students with ADHD/SLD in undergraduate STEM courses. Our data reflects the perception of our participants and does not include instructor interviews or classroom observations. We note that other studies have found student reports of active learning to match instructor reports of active learning (Andrews et al., 2011). We caution that additional research is needed to make conclusive statements about the efficacy of any one active-learning practice for students. However, many existing reports regarding active learning do not incorporate the voices of students with disabilities. Data were collected only at one institution. Limiting our data collection to one institution benefitted our analysis because we held more extensive background knowledge of instructors known to use active learning in their courses and the history of active learning on campus. All of our participants were currently registered with the DRC. It is unclear if our research findings apply to students who qualify for accommodations but are not yet receiving formal accommodations. A majority of our participants were white, men, and were majoring in the life sciences. We attempted to address this limitation by presenting quotes from all participants so that additional viewpoints are represented in our findings. More research is needed to determine if our results apply more broadly to students with other types of disabilities besides ADHD/SLD and to address how students with disabilities from more diverse gender and racial backgrounds perceive active learning to influence their learning and self-advocacy experiences. We anticipate that additional supports and barriers may exist.

## Results

We interviewed 25 STEM majors with ADHD/SLD to determine how the implementation of active-learning practices affected their perceived learning experiences in undergraduate STEM courses. Throughout the Results we present both a figure and a table to portray similar information. We are providing these options in an effort to align with principles of universal design for learning (*See Discussion*). The goal of this type of presentation is to provide readers the option of selecting the format which best suits their needs or preferences. Tables may be more accessible than a figure for people using a screen reader, while other people may prefer a figure. Participant quotes have been lightly edited for brevity and clarity. Brackets represent added text to enhance readability, and ellipses indicate text removed from the quote.

### Participants perceive themselves to learn best in a variety of contexts

During the interview we asked participants, “Do you learn better in a STEM course that uses active learning or in a STEM course that uses lecture?” Twenty-four of our 25 participants reported a preferred learning environment (Figure 1). One participant, Olen, expressed no preference, and another participant, Jessa, explained that she perceives herself to learn best outside the class regardless of if the STEM course uses lecture or active learning. Three participants expressed a preference for lecture-only courses. Seven participants shared that they preferred to learn in a course incorporating a mixture of lecture and active learning. Three participants stated that they prefer active-learning courses in a conditional manner. We considered these participants to represent a group called, “conditional active learning.” Participants in this group stated that they preferred active learning as long as they were able to sufficiently prepare for the course, as long as the instructor provided sufficient teaching materials to help them prepare outside of class, or if they were taking a STEM course in a particular STEM discipline (e.g., they preferred active learning for an engineering course, but lecture for a math course). Most of our participants (n=10) explained that they perceived themselves to learn best in an active-learning STEM course with non-conditional statements about this preference. Figure 1 summarizes the type of learning environment participants perceived themselves to learn best in, and Table 3 summarizes the figure while providing representative participant quotes.

**Figure 1.**
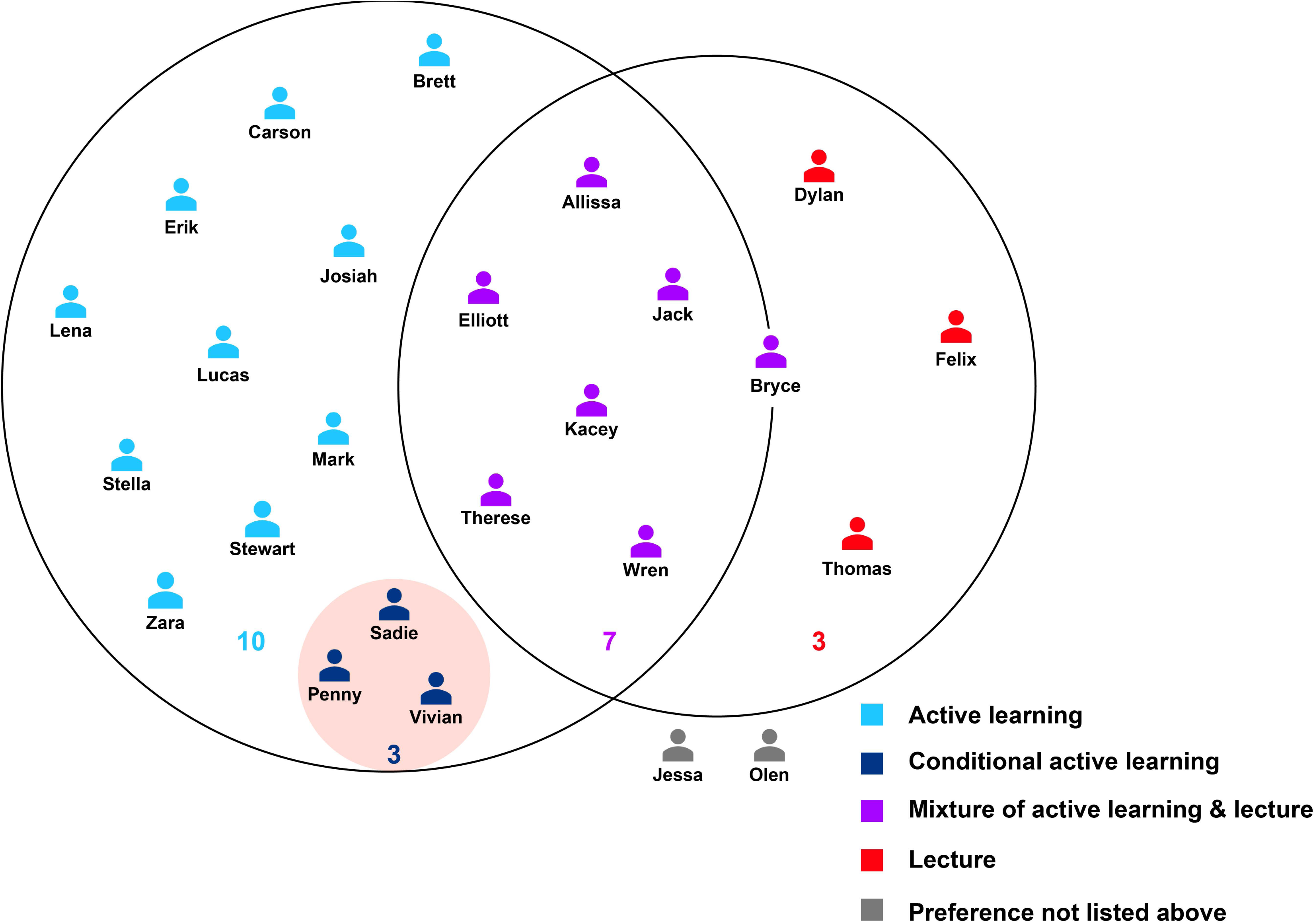
Participants perceive themselves to learn best in a variety of learning environments. Light blue indicates active-learning only courses, purple indicates courses with a mixture of active learning and lecture, red indicates lecture-only courses. Dark blue in a small red circle represents “conditional active learning.” Participants in the conditional active learning category described themselves as learning best in an active learning course if a particular condition was met (e.g., If I am prepared, I learn best in an active-learning STEM course). Gray signifies participants who perceived themselves to learn best in an environment not listed above, or who did not express a preference.

**Table 3.**
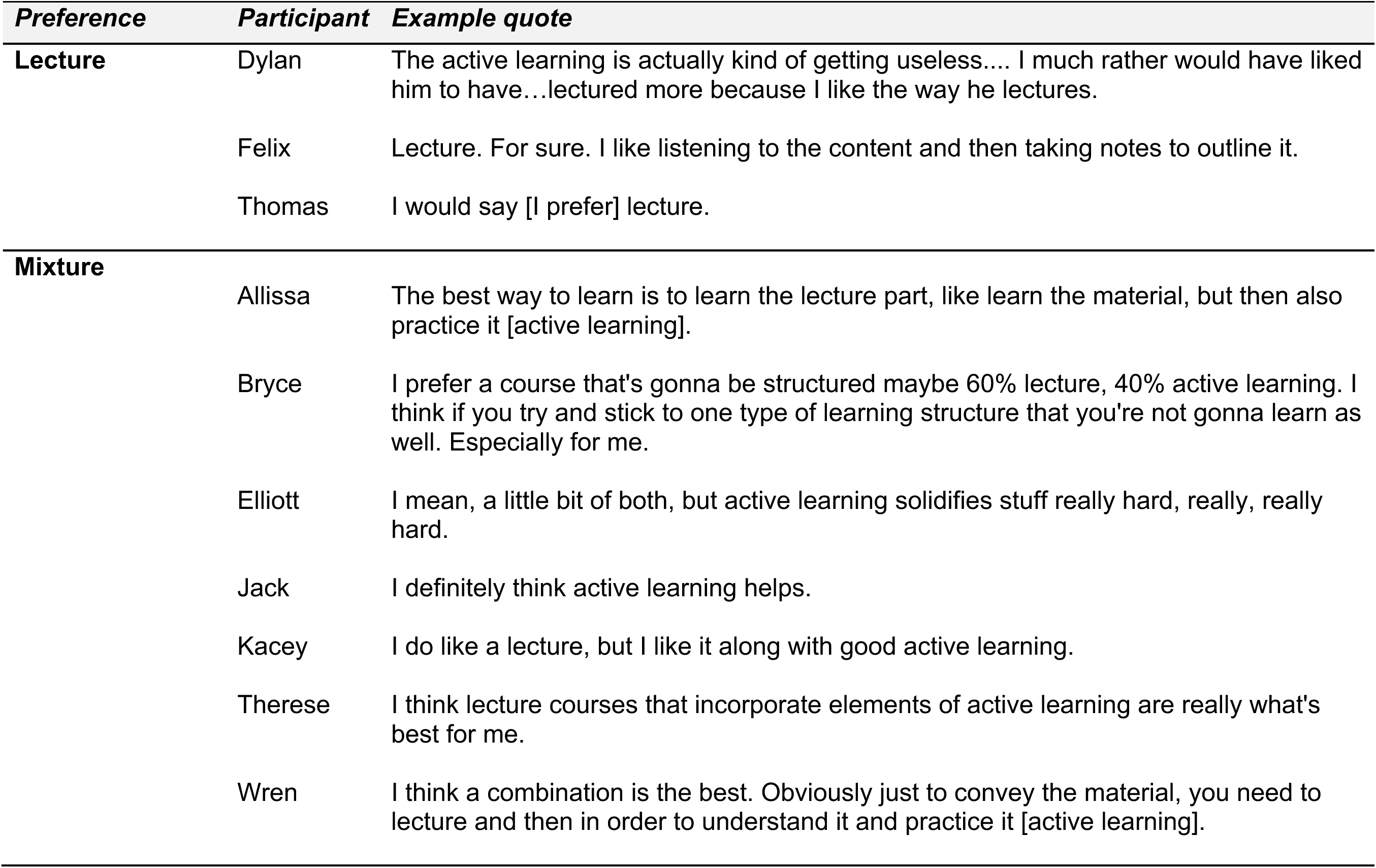
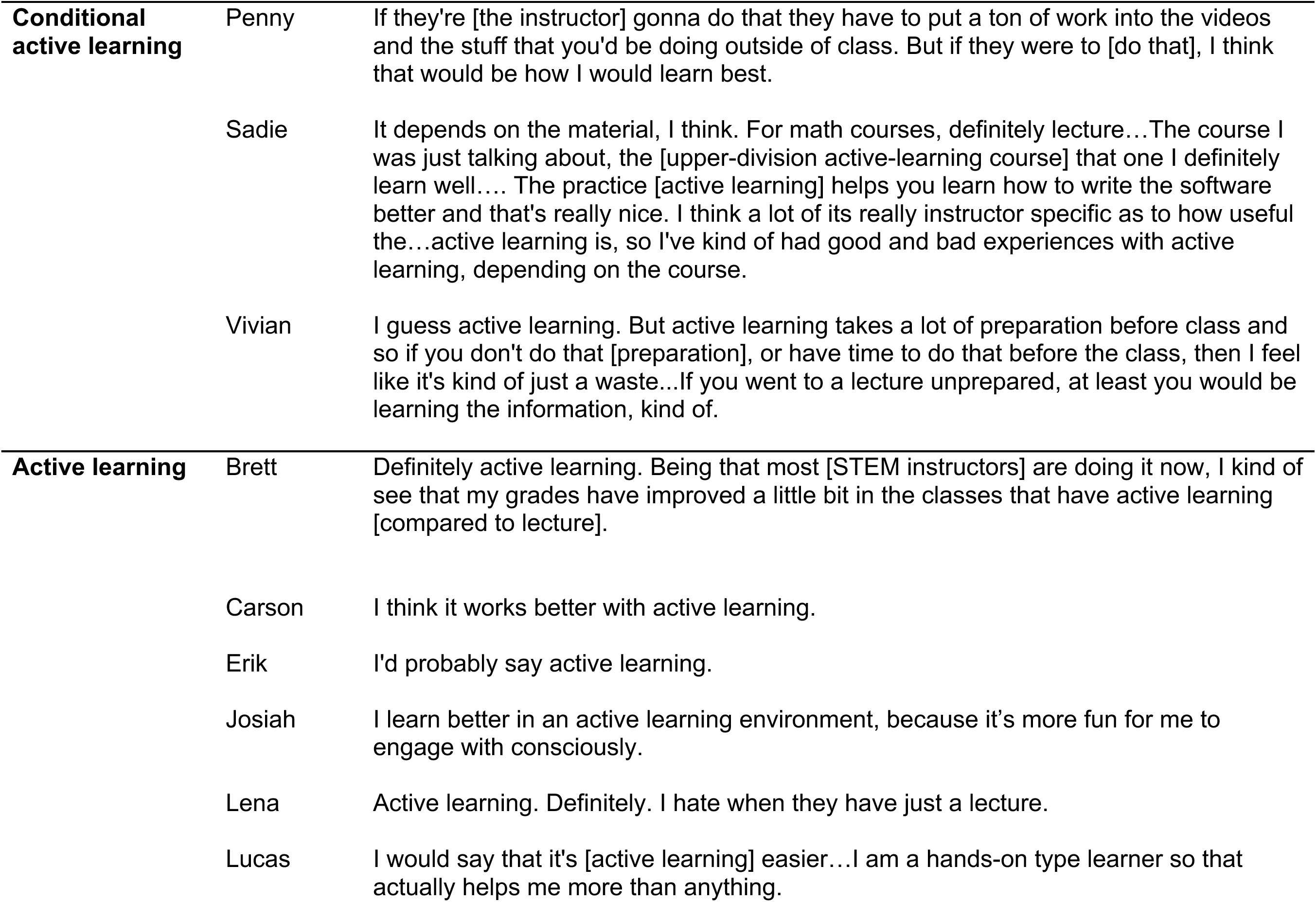
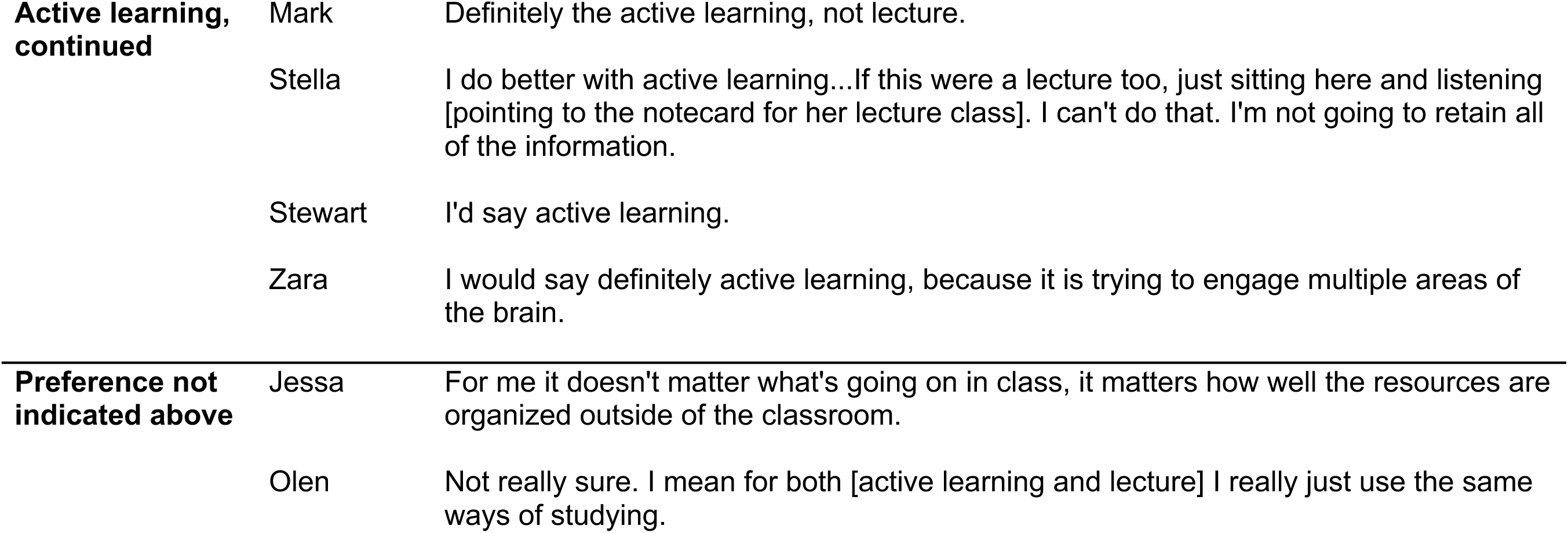
Participant responses to the question, “Do you learn better in a STEM course that uses lecture or active learning?”

### Aspects of active learning influence participant perceptions of learning

Participants described aspects of active learning that influenced their perceptions of learning in response to open-ended interview prompts regarding their active-learning STEM course. These interview prompts included: (1) Walk me through what a typical day is like for you in your active-learning STEM course, (2) Tell me about your interactions with your instructor, and (3) Tell me about your interactions with your peers. From our analysis we found several aspects of active learning which affected our participant’s perceptions of learning. Influential aspects of active learning included: environment, course structure, instructor reveals thinking, course materials, flipped courses, group work, and clickers. We begin by presenting the more general aspects of active learning which influenced our participants’ perceptions of learning (i.e., environment and course structure) and then move into more specific aspects of active learning, for instance, group work and clickers. Aspects influencing participant perception of learning are summarized in Table 4 (*See Supplemental File 4 for figure)*.

**Table 4.** Aspects of active learning influencing participant’s perceptions of learning. ^a^ The supports for flipped courses overlap with all other aspects of active learning functioning as a support. A few key supports for flipped courses are indicated in the table.

| ***Aspect of active learning*** | ***Supporting perceptions of learning*** | ***Hindering perceptions of learning*** |
| --- | --- | --- |
| **Clickers** | Promote metacognition | Short answer time limits<br>Distracted by class short answers |
| **Group work** | Promotes metacognition<br>Alternative sources of motivation<br>Builds relationships to ask questions and form study groups | Peers don't want to work with me<br>Potential for negative emotions<br>Lack of clear directions |
| **Flipped course <sup>a</sup>** | Flexibility in use<br>Promotes metacognition<br>Incentivizes attendance<br>Breaks up lecture | Long videos<br>Misalignment of video to class<br>Lack of instructor explanation<br>Withdrawing from course |
| **Course materials** | Organization of content<br>Flexibility in use | Extensive textbook reading<br>Lack of screen reader |
| **Instructor reveals thinking** | Promotes metacognition<br>Helps students apply content from worked examples | Lack of instructor explanation |
| **Course structure** | Incentivizes preparation<br>Incentivizes attendance<br>Frequent opportunities for practice | Overlooking accommodations<br>Increased stress<br>Challenging to prepare for |
| **Environment** | Encourages participation<br>Breaks up lecture<br>Opportunities for hands-on learning<br>Space for distractions | Room is distracting<br>Class moves too fast<br>Class moves too slow |

A definition of each aspect of active learning and a description of how the aspect influenced perceptions of learning is described in the following subsections. Within each subsection, we first demonstrate how the aspect of active learning supported our participant’s perceptions of learning. We then describe how this aspect hindered their perceptions of learning. Several aspects are overlapping in nature. The data presented within one aspect also applies to a different, but related, aspect of active learning. We present these aspects separately so that readers may readily identify a certain active learning aspect of interest and its associated supports and barriers to participant perceptions of learning.

#### Environment

Environment encompassed the physical space of an active-learning STEM course and the way a participant perceived the classroom climate of the course. When functioning as a perceived support of learning, the environment encouraged participants to participate during the class, broke up lecture into more manageable blocks of information, offered opportunities for hands-on learning, and provided space to be distracted in class. Participants like Wren, Elliott, and Lena shared that they feel comfortable answering and asking questions during class within active-learning STEM courses. Additionally, several participants reported that active learning divided up lecture and offered them other ways to engage with material besides listening and taking notes, which supported their learning. Dylan and Lucas were two participants with ADHD who especially valued active learning that provided opportunities for hands-on learning. Although Dylan perceives himself to learn best in a lecture-only STEM course he described in-class activities as helpful when they allowed him to physically manipulate objects with his hands as opposed to completing worksheets on paper.

> *Some activities can help, like the first activity helped out a lot, it was between primary, secondary, tertiary, and quaternary protein structure, and it was with phone cords. That helped a lot. But other [activities] can be challenging and then after you’re kind of just like, what was the point of that?… The ones that are good, I liked the phone cords because I had it in my hand. —Dylan*

Like Dylan, Lucas appreciated active learning because he considers himself to be a “hands-on learner.” He reported,

> *I’m a hands-on learner. I do better if I can see it and I do it. Some of the things that we were doing, I had done 100 times, but it didn’t help as far as actual classwork, but just getting the data myself and understanding like where it came from and how it was being used made me understand stuff so much better. — Lucas*

Finally, participants like Brett, described that the environment of active-learning STEM courses supports their learning because there is built-in space for them to be distracted without missing vital information. Brett stated:

> *With my ADHD, it’s great to kind of be able to get some energy out. Distractions are there for everyone and you kind of can work through them easier than in a lecture where if you get distracted you miss stuff. So active learning is definitely a huge benefit for any STEM class. —Brett*

While many participants found the environment of active-learning STEM courses to support their learning, there were ways in which the environment hindered their perceptions of learning. Stella shared that the multiple dry-erase boards present in a SCALE-UP-style room make it challenging for her to follow the instructor’s explanation.

> *The classroom setup is strange and irritating. There’s a lot of big, round tables around the room. There’s [dry-erase] boards here, here, and here and then a random teacher desk, tall desk in the middle with a computer …It’s irritating because he’ll be on one end of the room starting a problem, run over to the other end of the room, finish the problem… Half of the class has to move to see what he’s writing. There’s that dumb desk in the middle with the computer and everything. He’s just running back and forth. For me, with ADHD and stuff, it’s better to just have it all in one spot. So I’m not missing half of what you’re saying. —Stella*

Besides the physical layout of the classroom, participants reported that the pacing of their active-learning STEM course could hinder their learning. Both Brett and Allissa, who receive accommodations for ADHD, shared that often the in-class activities move too fast. Allissa said, *“When we do active learning in class, a lot of time it’s moving at the pace of the rest of the class so I’m like, “Oh crap. I haven’t finished yet.”* One participant, Dylan, felt that in-class activities moved too slowly. This suggests that unless the activity involved hands-on opportunities, he would often lose interest in completing the task.

#### Course structure

Another general aspect of active learning represented in our data was course structure. Drawing on prior work, we considered how participants perceived course structure (Eddy and Hogan, 2014). Specifically, our participants discussed graded preparatory assignments (e.g., frequent reading quizzes) and in-class engagement activities (e.g., clicker questions, worksheets, case studies) during the interviews. Some examples of in-class engagement, such as clicker questions and group work, are presented in our study as stand-alone themes due to the prominence of these aspects in our data.

We found that course structure incentivized class preparation and attendance. Jessa, a participant with ADHD, who perceived herself to learn best outside the classroom shared her experience with pop reading quizzes in an active-learning STEM course.

> *What was special about this class is that he would do pop quizzes at the beginning of class for attendance…I kind of hated them in the beginning, but they forced me to stay on track with the material and it was great. —Jessa*

Jessa explained that the course structure in this class encouraged her to prepare for each class, instead of learning content in fewer, more extended sessions. Allissa, who preferred to learn in a course with both lecture and active learning expounded on how course structure supported her learning in a STEM course because the frequent opportunities to practice the material in class prepared her for the exam.

> *I think a lot of concepts in STEM courses are things like problems and problem-solving things… I feel like practice is almost the most important part of learning the material, to be able to practice it for the exam. —Allissa*

Allissa valued the opportunity to practice problems in class so much that she reported that she had withdrawn recently from a lecture-only STEM course because opportunities to practice the problems in class were not provided.

Although participants found course structure to support their learning, there were ways in which course structure hindered participants’ perceptions of learning. Due to the increased frequency of assignments and quizzes, several participants explained having to engage in self-advocacy to ask their instructor to apply their extended-time accommodations to these assignments, which we refer to as overlooking accommodations. Kacey explained an ongoing situation she recently encountered in an active-learning STEM course using graded pre-class reading quizzes,

> *I know she has so many students, but she’s not being mindful of the ones with accommodations. She hasn’t submitted an agreement, and she’s not giving me extra time that’s already set up. That would normally already be set up on quizzes…. I just don’t know if maybe because they’re just like pre-class quizzes no one gets extra time, or if she just doesn’t realize that she needs to add it on. I don’t know. —Kacey*

At the institution where data were collected, instructors are sent an official accommodation notification through an online system. Instructors are asked to acknowledge that they have read these accommodation notifications, which is what Kacey refers to as “an agreement” in her quote. Kacey shared that she is in the process of addressing this reading quiz issue with her instructor.

Jessa was another participant who reported asking her instructor for extra time on pop quizzes in her course. She also stated that she found her active-learning STEM courses to be stressful, presumably due to the number of assignments and quizzes she was responsible for completing. Her feelings of stress could become elevated to such a degree that Jessa would decide not to take her ADHD medication on some days. Some of our participants discussed how taking their ADHD medication causes them undesired side effects. The side effects discussed by our participants included sleep problems, decreased appetite, and increased nervousness, which can affect their experiences in the classroom. Jessa stated:

> *I don’t like active learning. I kind of hate it when they put me in there…It really bothers me…It was a stressful class. It was stressful, especially because I’m not one that likes to go to class and to remember to take a pill at the beginning of the day, sometimes I don’t like taking it ’cause it’ll stress me out. —Jessa*

Many participants also found it challenging to prepare for assignments and class engagement activities. Often these challenges were related to issues with course materials and flipped courses (described below).

#### Instructor reveals thinking

Instructor reveals thinking encompassed instances when participants discussed the instructor or teaching assistants demonstrating or explaining course content. They often revealed their expert thinking through worked examples, after clicker questions, or by answering participant questions individually during class time. Felix, a participant with both ADHD and SLD, preferred to learn in lecture-only courses, described only one of his active-learning STEM courses as “acceptable.” He explained that the reason he found this particular active-learning STEM course “acceptable” was because:

> *At the beginning of every class, she would go over one problem, very in depth. She would try to pick the harder problems that she was giving that week to go over in class on the board. Then after the time that she worked the problem, we would have the rest of the class to ask her questions or ask the TA questions. —Felix*

When the instructor completed difficult worked examples, followed by the opportunity to apply this new information to example problems in class, Felix felt active learning supported his acquisition of knowledge. Many participants also expressed how the instructor revealing their thinking after closing a clicker question supported their perceptions of learning.

> *She’ll pose a question to us, we’ll work through it and then submit it…Afterwards, she’ll basically work the question out in full so that anyone who got it wrong knows how to do it, and then anyone who got it right reinforces that they did it correctly, and things like that. Works really well*. —Bryce

As illustrated by Bryce’s quote, the instructor revealing thinking overlaps with other aspects of active-learning, specifically clicker questions in this example. Our data further revealed that a lack of instructor explanation, or when the instructor failed to reveal their own thinking, had a negative impact on participant’s perceptions of learning. This view was most prominent for Kacey who shared her frustration regarding an active-learning STEM course. She explained that her instructor failed to explain why certain clicker question answers were correct or incorrect. Kacey stated:

> *I don’t understand how you could call this a class, when they’re just throwing this stuff at you, but they’re not helping you understand it, and they’re not going back and saying, “This is wrong because of this, and this is how you do it the right way, because X, Y and Z.” I got none of that from her, and so I learned all my stuff from [a third-party tutoring service]. –Kacey*

Kacey’s dissatisfaction when the instructor failed to reveal their own thinking was shared with Thomas. Thomas stated that when his instructor does not explain the content or the purpose of an activity before starting group work, he doesn’t understand “what the hell’s going on.” This made active learning challenging because Thomas was unsure of what he was supposed to learn and what he needed to accomplish during the class period. In this type of situation a student may feel lost, or incapable, because they do not understand the expectations and objectives for the class period, which can make it challenging to communicate with their group mates. These issues can, in turn, lead to the student being excluded from the group.

#### Course materials

Course materials comprised of media and other tools, such as notetaking supports, provided by the instructor with the presumed purpose of supporting student learning of class content. Participants discussed how course materials such as videos and notetaking guides supported their learning by promoting organization of content and flexibility in use by the student. Lena, a participant with ADHD, explained how the videos provided by the instructor supporter her learning in a STEM course.

> *I’ll be watching it, but I have to rewind because I’m not paying attention, so I like the fact that because you can’t do that in real life. I have to rewind things and that really helped. The class was still hard, but I think I would have failed that class if that wasn’t a thing. Because I could just rewind it as much as I can and write. Pick up on things and if I had questions, I could write them down. Ask the professor later in [class]. –Lena*

Another participant, Wren, who qualifies for a notetaking accommodation due to ADHD, described that the interactive notetaking guide provided by her Calculus instructor supported her learning. The interactive notetaking guide was helpful for her because it supported organization of class material and created a resource that she could access while solving practice problems.

> *We were actively engaged when he was lecturing. We had the packet to refer to when we were [solving problems] that were like notes, but notes that made sense and we didn’t have to waste time writing down, like, oh, it’s this theory… [The notes] made a lot more sense. I really excelled when I had that type of resource. —Wren*

While course materials supported the learning of many participants, other participants explained that some course materials were unsupportive of their learning. A common issue was difficulty reading from the textbook, which affected a participant’s ability to prepare for in-class activities. Vivian, who has an SLD, expounded on this by saying:

> *It’s hard for me to read something without being exposed to it before, I guess, so if I just read about a topic, then I really don’t grasp it at all. I prefer to read after the class, so in classes that they don’t want you to do that, it’s really hard for me*. –Vivian

Vivian and other participants explained that when the textbook and other materials are inaccessible to them, active learning can be a very negative experience. Reading materials may be inaccessible to students when a student qualifies for an audio version of their textbook, but they do not have or are not able to use this technology for a variety of reasons. We also considered the textbook to be inaccessible to students when the sheer volume (e.g., 80-100 pages) of required reading for a single class period is not feasible for them to complete when taking multiple classes at the same time in a short window of time. Many of the supports and barriers associated with course materials were related to flipped courses.

#### Flipped courses

A flipped course is a type of class in which students are introduced to material outside the formal class period before coming to class. During class, opportunities for applying and practicing the materials are provided that often involve additional active-learning practices (e.g., group work, clickers, etc.). Flipped courses are considered by some researchers to be a form of active learning and were discussed by many participants as an active-learning practice in the interview (D. Lombardi & Shipley, 2021). We found that flipped courses supported participant perceptions of learning in the same ways described for other aspects. That is, the supports from environment, course structure, instructor reveals thinking, course materials, group work, and clickers applied to flipped courses. However, we present flipped courses as a stand-alone aspect of active learning because flipped courses represented the most profound active-learning barrier reported by our participants. Many of our participants perceived that they were expected by their instructors to learn material individually, as opposed to just becoming familiar with terminology or basic concepts outside of class, before the next session. Two participants, Bryce and Penny, reported deciding to withdraw from a flipped course, which ultimately altered their plans to complete their degrees. Bryce, a student who qualifies for alternative textbooks (an audio version of the textbook) explained that at this particular time he was not yet able to access his accommodations because he was in the process of being re-evaluated in order to provide official documentation of his disability to the university disability resource center.

> *The reason I [withdrew from a flipped STEM course] was hugely in part due to the “flipped” classroom setting. Basically, it was 80-100 pages of reading in between classes in the textbook. And, that was, essentially, how you were supposed to teach yourself the course. In terms of what we actually did in class, I was amazed. There was no lecture component to the course whatsoever. You would walk into the course, you would sit down, and then immediately at the start of the course, you would start answering the clicker questions. And, after each question, there was no explanation of why that was the answer was this, that, or the other. It was just, “Let’s move on to the next question.”… When I was doing the reading, I wasn’t pulling all the information in, just because there was so much of it. And, therefore, I was struggling on these questions in class. And, I had no lecture, no component to it where I was getting taught the information. I struggled so much that I felt there was no way I was gonna succeed. And, that was the main reason I dropped out. So, I’m not looking forward to retaking that course. —Bryce*

Bryce’s negative experience with flipped courses was reiterated by Penny. Penny was a participant with both ADHD and SLD who stated,

> *I withdrew from [a flipped STEM course] because it was really bad. [The instructor’s] videos that he would make were really unhelpful. He would give very simple examples and then in class he would give us extremely hard problems, and they just weren’t helpful at all. –Penny*

Other participants described difficulties in flipped courses, as well. Lena shared that at first it was challenging for her to learn how to prepare for her flipped STEM course.

> *It was kind of weird adjustment at first, because I’m not used to getting YouTube, like videos sent to me, so it was kind of like you had to pace yourself and watch it right at the right time. —Lena*

Lena explained that she can struggle to schedule and plan her work as part of her disability. Felix, a participant with ADHD and an SLD, explained that he struggled to pay attention to videos. Felix explained that, *“When the video lectures that you watch at home aren’t very engaging, I do not pay attention to them at all. I just can’t make it through them.”* Later in the interview Felix shared that because he struggles to watch the videos, he comes to class unprepared, which has a negative impact on his experiences in group work.

> *[The thing] I didn’t really like so much about the flipped classroom thing was if you had fallen behind on lectures or homework, it just made me feel uncomfortable to struggle through questions working with other people. –Felix*

Felix’s data is one example of how issues with flipped courses relate to other active-learning practices. In this case, the related aspect of active learning is group work.

#### Group work

Group work was an aspect of active learning discussed by participants during the interview. We defined group work as instances students described working with their peers to complete a task or assignment. Group work supported participant perceptions of learning by providing feedback, which supported metacognition. This was illustrated by Allissa when she said, “*When you discuss with your peers in class, you’ll find out right then, okay this is where I’ve been thinking about this wrong.”* Here, Allissa alluded to how she uses group work to monitor her own understanding of class concepts in the moment. Elliott similarly described how group work supported his metacognition:

> *It helps you learn when you have to argue your point against a group of people and you know you’re right, or you might not be right but you have to argue your point, and by arguing your point you have to prove it to them, and then in proving it you find out whether you’re right or you’re wrong. I guess it’s helpful because it helps you make those connections. —Elliott*

Group work further supported participant perceptions of learning by providing alternative sources of instruction and motivation for learning. Several participants stated that they found group work valuable because their peers would explain concepts at a level they could more easily understand compared to how their instructor explained concepts. Carson, an upper-division student with both ADHD and SLD, shared that he no longer feels motivated to learn in his courses by his own grades. He explained how group work provided him with a different motivation to learn.

> *I think [group work] engages me more, personally, because I work best when I’m helping other people. Because if it’s just like about me, and like my grade, and like that kind of stuff it’s like I don’t know. I have a hard time motivating myself to like get straight As. There’s like I guess a bunch of different reasons for that. But like, if I have to work with someone on a problem, then it feels more real, feels like reality… It motivates me to like learn the material, because I enjoy explaining things that I understand. –Carson*

Group work also supported our participant perceptions of learning by fostering peer-to-peer connections. Participants described that they could ask their peers questions about topics they found challenging in the future, and that they could form study groups because of group work in class. Peer-to-peer connections are also likely a support because it provides a mechanism to fill gaps, a self-advocacy behavior.

Several components related to group work were perceived to hinder the learning of our participants. These barriers included instances when peers did not want to work with the participant because the participant was thought to work more slowly than the class. Lena, who is an upper-division Engineering student, stated:

> *Engineers tend to be very impatient. So if you’re not up to speed with them, they’ll just not work with you anymore. And I guess that’s understandable, I guess you don’t want anybody to hold you back, but if they did [work with me], active learning would probably be ten times better. —Lena*

Other participants explained that group work can create the potential for them to experience negative emotions during class. Jack, a student with SLD in reading, reported:

> *I wish they would understand why I never like reading in groups like out loud reading or why I don’t like writing by hand in front of them. I could always tell them, “Don’t feel like it,” most of the time, because it’s just some random group I have to work with. But if they knew why I didn’t want to read out loud or why I didn’t want us to write then they probably wouldn’t make me, unless they were just really lazy and they didn’t want to do it themselves. But I wish they would know. If they knew why I was struggling, it’d be better than them just thinking I was an idiot. –Jack*

Other participants like Erik shared this sentiment. Erik explained that as someone with an SLD in writing, he feels especially uncomfortable being asked to hand-draw graphs in front of his peers. Moreover, group work can be a time in which participants will have to reveal their use of accommodations or discuss their disability with their peers. Zara shared that she discussed why she uses extended time accommodations when her peers noticed she received extra time for quizzes.

> *It was brought up, because we did group quizzes and I got longer time on a group quiz and they go like, “Why do you get longer time?”—Zara*

Zara was a participant who felt comfortable explaining why she used accommodations to her peers. However, many of our participants were not comfortable talking about their disability or accommodations with their peers. We anticipate that if these participants were in a group exam situation, similar to what Zara described, they would feel highly uncomfortable. Lastly, participants reported that a lack of clear directions or expectations for group work from the instructor hinders their perceptions of learning. Participants, like Stewart, shared that they feel frustrated when the instructor does not provide clear directions for group work.

#### Clickers

Clickers are student response systems that allow students to anonymously share their answers to instructor questions. Answering clicker questions supported the metacognition of our participants by prompting them to monitor their own understanding of class content. Participants described especially valuing the instructor’s explanation of why each answer option was correct or incorrect. However, some implementation of clicker questions within STEM courses was problematic for participants, notably when clicker question responses required them to compose short answers within a limited amount of time. Josiah, a participant with SLD in reading, and Stewart, a participant with ADHD, explained that they felt they needed more time to type their answers than provided by the instructor. This was especially concerning for Stewart because in one of his classes his clicker questions were graded for accuracy. Other participants, like Kacey, explained that viewing the class responses to free-response clicker questions is particularly stressful for her due to differences in formatting.

> *With [free response clicker questions] the technology is weird, because it’s like you either type in ABCD in all caps, or you type in your answers that you think are all right, so if all of them are right, you would type them all in. Some people do them in all capital letters, some people do them in all lowercase letters. Some people do them in both kind of letters. Some people put in commas, some people don’t. That’s so stressful for me. It’s not organized in that way. —Kacey*

Some participants with ADHD shared that the way information is organized affects their learning because they will attend to details their instructor does not consider to be important, but these non-essential details seem and feel very important to the participant. This results in situations where the participant is spending time making sense of these non-essential details on their own, which can detract from learning what their instructors consider to be essential details leading to stress. Kacey and other participants expressed a strong preference for clicker questions with multiple-choice responses that are aggregated automatically by the clicker’s software program. They found this format less distracting so they could focus on the content of the question and the submitted responses as opposed to the formatting differences of the answers submitted by the entire class.

### The influence of active learning on self-advocacy

We asked participants if their self-advocacy changed as a result of being enrolled in a STEM course that incorporated aspects of active learning. Participants shared a diverse array of responses to this question, which are summarized in Figure 2 and Supplemental File 5. Broadly, there were two main groups of participant views. Participants who did not think their self-advocacy changed, and participants who thought their self-advocacy changed in response to aspects of active learning. For participants who reported that their self-advocacy did not change, there were two distinct reasons. The first reason involved how the participant viewed their accommodations. For these participants, they reported their self-advocacy did not change as a result of active learning because their accommodations should always be sufficient for them. This view was illustrated by Sadie who said:

**Figure 2.**
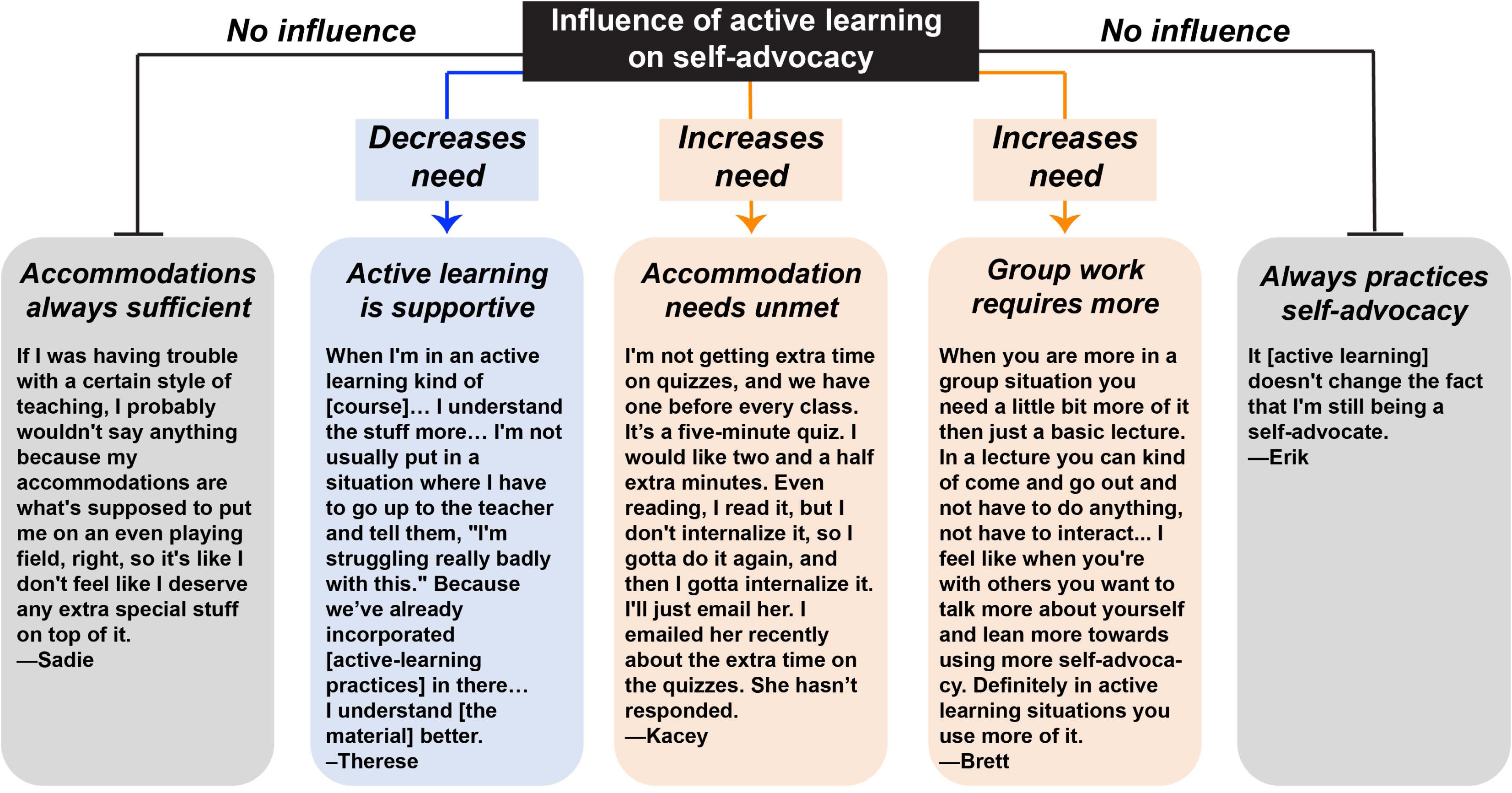
The influence of active learning on self-advocacy. Participants explained that they saw active learning as exerting no influence (gray) on their self-advocacy, or that they considered active learning to influence their self-advocacy (blue and orange). Blue represents a decreased need for self-advocacy, while orange represents an increased need for self-advocacy.

> *If I was having trouble with a certain style of teaching, I probably wouldn’t say anything because my accommodations are what’s supposed to put me on an even playing field, right, so it’s like I don’t feel like I deserve any extra special stuff on top of it. —Sadie*

Here, Sadie’s notion of self-advocacy is equated with only the use of accommodations. She did not consider it appropriate for her to communicate with the instructor if there are aspects of the course that may not support her learning. Other participants shared that active learning does not influence their self-advocacy because they are positioned to self-advocate in all course types. In other words, these participants would engage in self-advocacy at the same level in all their courses regardless of if it was an active-learning or a lecture course. Erik articulated this view. “*It [active learning] doesn’t change the fact that I’m still being a self-advocate.”*

Although some participants did not view aspects of active learning to influence their self-advocacy, many participants did describe an influence. A handful of participants stated that they perceive aspects of active learning to decrease their need for self-advocacy. These participants tended to find active learning to be supportive of their learning. For example, Therese, who prefers to learn in a STEM course that incorporates both aspects of active learning and lecture, stated.

> *When I’m in an active learning kind of [course]… I understand the stuff more… I’m not usually put in a situation where I have to go up to the teacher and tell them, “I’m struggling really badly with this.” Because we’ve already incorporated [active-learning practices] in there…I understand [the material] better. –Therese*

Therese saw active learning as providing her with more opportunities to learn the course material as opposed to lecture-only STEM courses. She also saw engaging in self-advocacy as a means to support her learning of course content. Because she feels her learning is better supported in a STEM course incorporating aspects of active learning, she perceived her self-advocacy needs to decrease. Yet some participants described that their need for self-advocacy increases as a result of active learning.

Some participants reported that their need for self-advocacy increases largely due to the issue of overlooking accommodations, which we described in the course structure section above. That is, with the increased number of assessments related to course structure there are more opportunities for the instructor to overlook their extra-time accommodations. In our study, this often occurred for pre-class reading quizzes, pop quizzes administered in class, and when clicker questions required short answers and were graded for accuracy. Kacey shared:

> *I’m not getting extra time on quizzes, and we have one before every class. It’s a five-minute quiz. I would like two and a half extra minutes. Even reading, I read it, but I don’t internalize it, so I gotta do it again, and then I gotta internalize it. I’ll just email her. I emailed her recently about the extra time on the quizzes. She hasn’t responded. —Kacey*

Other participants described that because of group work, an aspect of active learning, they perceive their need for self-advocacy to increase. Brett, a participant with ADHD, expanded on this notion.

> *When you are more in a group situation you need a little bit more [self-advocacy] then just a basic lecture. In a lecture you can kind of come and go out and not have to do anything, not have to interact… I feel like when you’re with others you want to talk more about yourself and lean more towards using more self-advocacy. Definitely in active learning situations you use more of it. —Brett*

While some participants perceived active learning to increase their need for self-advocacy, many of these participants explained that although there may be an increased need for self-advocacy they feel more comfortable communicating with the instructor. Communication is an essential component of self-advocacy (Pfeifer et al., 2020). Participants attributed enhanced comfort to communicate with the instructor to the environment of their STEM courses that incorporate active-learning practices. Stella reported,

> *When the teachers are walking around and stuff, that gives you the opportunity to ask more questions if you need it. It’s not all 100% the student going to the teacher, but more of this, this kind of equality. —Stella*

Lena shared a similar sentiment and noted that she feels more comfortable communicating with instructors in active-learning STEM courses because her peers are less likely to notice her talking to the instructor. Lena was a participant very concerned about the prospect of her peers learning she has a disability and qualifies for accommodations.

#### Examples of self-advocacy

During the interview, a handful of participants shared specific examples of self-advocacy in the context of an active-learning STEM course. Participants described asking the instructor to apply their testing accommodations to quizzes and clicker questions. Our participants also explained that they would fill gaps, a self-advocacy behavior, by asking peers from their group to explain challenging topics to them, and in some cases tutor them. Participants further filled gaps by seeking third-party tutoring when they found their active-learning STEM course did not support their knowledge acquisition. Moreover, participants utilized their self-advocacy knowledge and beliefs to make decisions about their own success and well-being in the course. The repercussions of these decisions have the potential for negative long-term effects. However, we consider these decisions to be our participants’ responses to an environment that was not designed with them in mind, and we do not fault them for these decisions.

At least one participant described that they decided to forego ADHD medication some days to mitigate feelings of anxiety in active-learning courses with frequent assessments. As described earlier, some of our participants experienced side effects from their ADHD medication that made them feel more nervous than they typically would if they did not take their ADHD medication. By deciding to forego ADHD medication, students may alleviate some of the stress they feel in an active-learning STEM course. But, foregoing ADHD medication frequently before class has the potential for negative long-term effects. Lucas, a participant diagnosed with ADHD in college, described what a STEM class was like when he was not taking ADHD medication.

> *It was a [STEM] class of probably about 60 or 70 people. [When other students were] like whispering, or talking, or tapping a pen, moving paper… Anything would just throw me off and I would miss a good bit of the [class]. And I found out that I was missing a good bit of the [class] ’cause… I recorded the class, I went back and listened to it, and I missed over 30 minutes of the class just from zoning out or wondering what somebody was talking about. –Lucas*

One of Lucas’s accommodations is to audio-record his classes. Later in the interview, he explained that when he takes his ADHD medication, he feels like he does not need to record class because he can pay attention throughout the duration of the class period. Other participants perceived that without ADHD medication their notetaking ability is diminished. However, we want to be clear that the decision to take ADHD medication (or not) is a complex and personal decision made by students. Our participants shared additional reasons, not directly related to the type of instruction in a STEM course, that influenced this decision.

Another decision made by participants included deciding not to read before active-learning STEM courses. Participants explained that because they struggled with the amount of reading and the unfamiliar vocabulary in their textbooks, they often felt like the effort was not worth it because they would not retain the information anyway. This decision had the potential for negative repercussions because when participants were unprepared for class, they described active-learning as “kind of a waste” and could feel uncomfortable working with others.

Finally, participants explained that they decided to withdraw from some flipped courses that hindered their perceptions of learning. In our interviews, we did not ask the participant if they talked with their instructors or their academic advisors before making this decision. Deciding to withdraw likely prevented the participants from receiving a low grade in the course. Yet withdrawing from a course has the potential for negative long-term effects because it can increase time to graduation if the course is required.

### Participant suggestions for STEM instructors about active learning

One of the final questions in the interview asked participants to share what they wished STEM instructors knew about their experiences in active-learning STEM courses. We organized their feedback into major points STEM instructors should be aware of when they incorporate active-learning practices into their courses. Quotes for each point are provided in Figure 3 and Supplemental Table 6. We encourage readers to reflect upon these quotes, which we used as the basis for the recommendations in the implications for teaching section in our discussion (Table 5).

**Figure 3.**
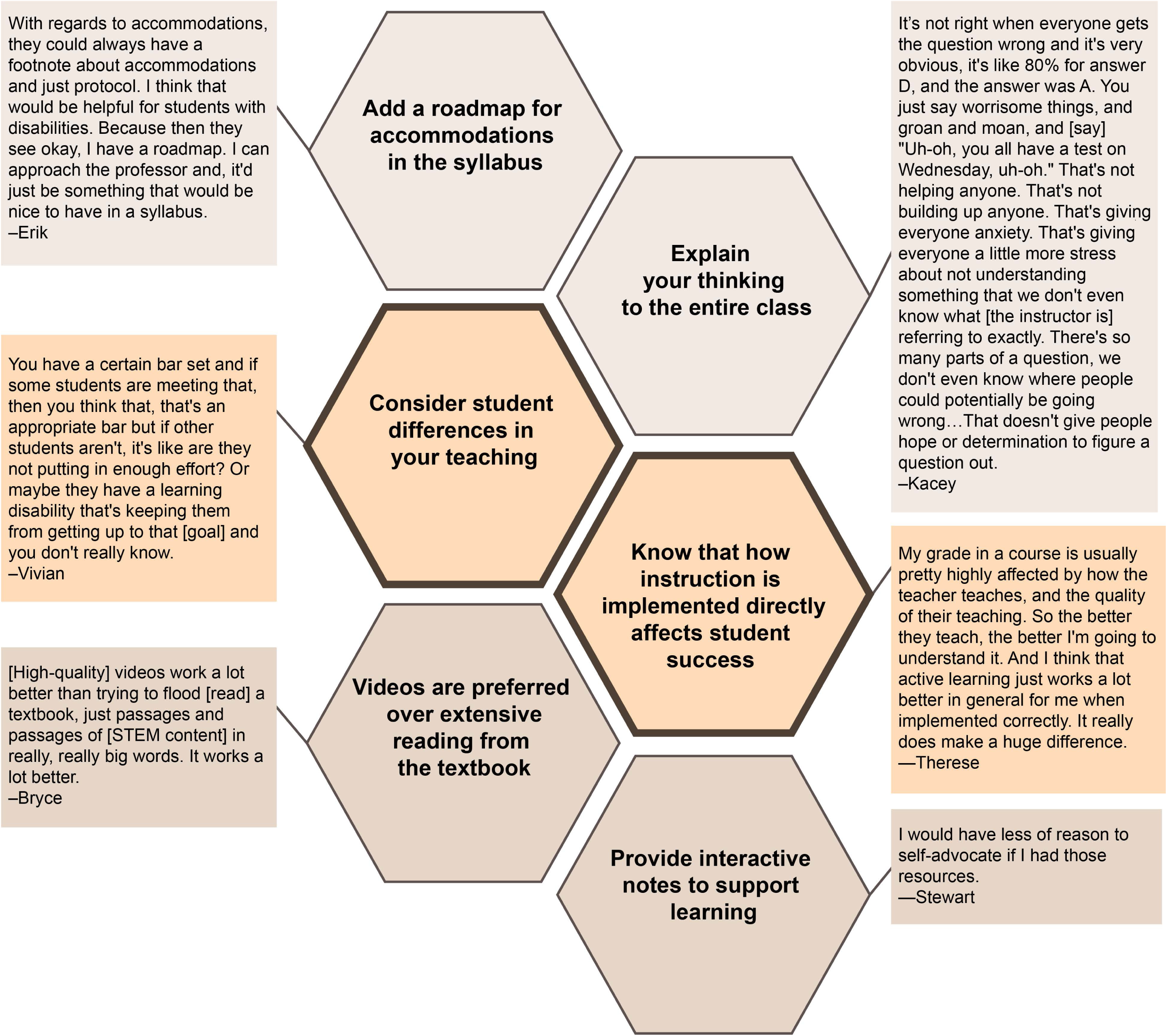
Participant suggestions for STEM instructors of active-learning STEM courses. Orange represents more general suggestions and shades of tan indicate more specific suggestions related to barriers presented in this study.

**Table 5.** Suggestions for STEM instructors. Bold text indicates suggestions offered by participants. We do not consider these suggestions to represent a panacea ensuring full accessibility of a course. These suggestions are founded on the specific barriers, and some of the supports identified by participants in our study.

| **Aspect of active learning** | **Suggestion** |
| --- | --- |
| **General** | **Consider student differences in your teaching.** Across our studies, participants shared that they wanted their STEM instructors to be more aware of how ADHD and SLD can affect their experiences in STEM courses (Pfeifer et al., 2020; Pfeifer et al., 2021). See Discussion for more information.<br>**Know that how instruction is implemented directly affects participant success in a course.** Several participants describe that active learning can be a significant support for their learning, if implemented appropriately. We encourage instructors to consult existing resources when incorporating active learning strategies into their courses, for example *CBE-LSE Evidence-Based Teaching Guides*.<br>**Add a roadmap for accommodations in the syllabus.** See Discussion for more description.<br>Conduct access check-ins regularly with your class to determine what students need in order to do their best work (Sins Invalid, 2019; Reinholz & Ridgway, 2021). Reinholz & Ridgway (2021) provide directions and several examples of how these types of check-ins can be incorporated into undergraduate STEM courses.<br>Review the checkpoints from the universal design for learning framework and incorporate them into the design of the course. As a starting point, we encourage instructors to review the guidelines and checkpoints within the principle called “providing multiple means of representation” (CAST, 2018).<br>Find ways to include “hands-on” learning opportunities for students when possible. For example, students can benefit from manipulating 3D printed models of structures that are difficult to understand. Participants in our study appeared to especially value these types of in-class engagement activities over more abstract, paper-based activities. |
| **Group work** | Provide clear expectations for group work and clear learning objectives for group assignments.<br>Offer options for students to opt out of a specific group role. For example, by offering an out, students who do not feel comfortable reading or writing in front of their peers could select a different role in their group.<br>Communicate the expectation that all group members should be included and establish a mechanism to ensure that all students are included in group work. This could look like frequent instructor check-ins to make sure students are included in their groups. Wilson et al., (2017) suggest that using reward structures (e.g., shared grades or certificates of recognition for reaching a specific goal) can incentivize students to work together. |
| **Clickers** | Avoid asking students to complete free-response clicker questions, especially with strict time limits, or as suggested by Gin et al., (2020) offer students the option to submit their responses before or after class.<br>Select clicker software programs that aggregate student responses. The volume of free-response text answers can be distracting to students because it can be challenging to focus on the content of the answers as opposed to the way the answers are formatted.<br>**Explain your thinking to the entire class.** Student learning is enhanced when students are provided the opportunity to discuss clicker responses with peers combined with instructor explanation of answers (Smith et al., 2011). |
| **Flipped courses** | **Videos are preferred over extensive reading from the textbook.** Use established evidence-based practices to create short, engaging videos that are closed captioned (Brame, 2016).<br>**Provide interactive notetaking guides.** Participants described that fill-in-the-blank style notes from the instructor supported their learning. This helped them take notes during the lecture portions of some STEM courses, and could also support textbook reading and video watching for flipped courses. |
| **Flipped courses, continued** | Organize video links and provide students with suggestions for how to use the videos to prepare for class. Be explicit about the length of the videos and invite students to take notes while watching. |
| **Course materials** | Select textbooks with built-in voice-to-text features that students can readily access.<br>Provide detailed reading assignment schedules to students, ideally by the first-day of the course. This helps students because they can share these schedules with the DRC to create accessible forms of readings in a timely manner. If you use primary literature or other reading sources not found in a textbook have PDF versions of these readings readily available. If you are contacted by the accessible media team at your Disability Resource Center you can provide the PDFs in a timely manner, which supports student access. |
| **Course structure** | Apply extended time accommodations to reading quizzes and graded clicker questions.<br>Offer students options to take pop quizzes before class starts or after class so they can use extended time accommodations without missing class instruction. |
| **Environment** | Invite students who feel highly-distracted in a SCALE-UP type classroom to meet with you to find the least distractable seat in the room. You could share this invitation verbally at the start of the class, or by posting it in your course syllabus or on the course website. |

## Discussion

In this study, we investigated how the implementation of active-learning practices in undergraduate STEM courses affected the perceived learning experiences of STEM majors with ADHD and/or SLD (ADHD/SLD). A majority of our participants perceived themselves to learn best in a STEM course incorporating at least some active-learning practices. Only three of our 25 participants reported that they perceived themselves to learn best in a lecture-only STEM course. This result is notable. Currently, much of the extant literature seems to suggest that active learning is most likely problematic for students with disabilities (e.g., Gin et al., 2020; Gonzalez, 2017). Here, we discuss how active-learning practices influenced our participant’s perception of learning by connecting our results to existing literature. We close by providing implications for research and theory, as well as teaching.

### The influence of active learning on perceptions of learning

Our results establish that there are enactments of active learning that our participants perceive to hinder their learning. We call these enactments active-learning barriers. Some of these active-learning barriers are previously characterized. Students with SLD in reading are known to experience difficulty with the amount of textbook reading required in college courses (e.g., Hadley, 2006). Additionally, SCALE-UP-inspired classrooms can be distracting for some students with ADHD (James et al., 2020). Interviews with DRC directors revealed that students with ADHD/SLD may not receive enough time to complete clicker questions, and that students with ADHD/SLD can experience difficulty working in groups (Gin et al., 2020). In our study, some students with ADHD/SLD described active-learning classes to be stressful, which tracks with existing active-learning research for students more broadly (Brigati et al., 2020; Cooper et al., 2018; Downing et al., 2020; England et al., 2017; Hood et al., 2021). While some of the barriers found in our study were previously known, we did identify a previously uncharacterized active-learning barrier for students with ADHD/SLD. This barrier involved how student responses to clicker questions are compiled for class discussion. When students submit short-answer responses to clicker questions, participants described being distracted and stressed by the formatting differences in the displayed answers.

There is clear evidence from our study that active-learning practices can be perceived to support participant learning. Our study and a study in introductory physics found that participants with ADHD perceived active-learning classes to provide “space for distractions” (James et al., 2020, p. 16). Many of the active-learning supports identified in our study are consistent with the recognized benefits of active learning. For example, we found that our participants appreciated course structure, e.g., frequent quizzes and graded assignments, because it incentivized preparation and attendance for class, and because it provided frequent opportunities for students to practice before the exam (e.g., Eddy & Hogan, 2014). Clicker questions and group work were valued because they informed our participant’s metacognition by helping them to monitor their own understanding of class content. Senior-level biology students reported that group work provides opportunities to monitor their own understanding of class content and group work can prompt students to explain their reasoning to themselves and their peers (Stanton et al., 2019; Wilson et al., 2017). Because we only interviewed students with ADHD/SLD, we did not determine how the supports named in our study compare to the supports that may be identified by students without ADHD/SLD. We hypothesize that some of the active-learning supports described in our study may be of greater importance for students with ADHD/SLD. For example, students with ADHD/SLD may be more likely than students without ADHD/SLD to report benefitting from well-designed videos in flipped courses because of the ability to re-watch videos as needed during their learning process. Future research is needed to determine if this and other similar hypotheses are supported.

In our study we found that a single active-learning practice could function as a support or as a barrier for our participants. From this finding, a question emerges. How can the same active-learning practice support learning in some cases, but hinder learning in other cases? We begin to address this question by discussing two instruction-related issues: implementation of an active-learning practice and universal design for learning. We first examine implementation.

### Implementation issues contribute to formation of active-learning barriers for students with ADHD/SLD

Many of the barriers reported by our participants appear to arise from how a particular active-learning practice is implemented. For example, both Bryce and Penny explained that they withdrew from flipped courses. In Bryce’s case, this flipped course included extensive readings from an inaccessible textbook and a lack of instructor explanation following clicker questions. Research regarding flipped courses shows that videos over textbook reading led to enhanced student performance (Jensen et al., 2018; Pulukuri & Abrams, 2021). For Penny, it appeared that the instructor’s videos were not following established best practices for the creation of educational videos (Brame, 2016). She shared that videos in this flipped class felt disconnected from what they were doing in class, which suggests that the videos were not created by the instructor with relevance to the course in mind (Brame, 2016). Our participants also described an active-learning barrier when the instructor fails to explain why clicker question responses are correct or incorrect. Previous research regarding the use of clicker questions shows that providing students with a combination of peer discussion and instructor explanation improves student performance (Smith et al., 2011). Overall, our data suggest that how an active-learning strategy is implemented, as opposed to the particular strategy itself, has the potential to impart severe academic consequences for students with ADHD/SLD. This is concerning, and something that as a community, we should take steps to address.

### Universal design for learning in an active-learning context

Based on our results, one way to enhance STEM active-learning experiences for students with ADHD/SLD is to consider using universal design for learning. Universal design for learning is a framework of three guiding principles, nine guidelines, and 31 checkpoints that build from general to specific suggestions for instructors (CAST, 2018). The three guiding principles of universal design for learning are to provide (1) multiple means of engagement, (2) multiple means of representation, and (3) multiple means of action and expression. Universal design for learning was originally developed to create more accessible classrooms for students with disabilities in K-12 classrooms (Jimenez et al., 2007). While further research of universal design for learning is needed to fully understand its utility in supporting the learning of all students in college courses, we see value in considering this framework and how it relates to our study (Boysen, 2021).

Previous studies of undergraduate STEM curriculum and courses show that the adoption of universal design for learning is limited (Scanlon, Legron-Rodriguez, et al., 2018; Scanlon, Schreffler, et al., 2018; Schreffler et al., 2019). Our results are consistent with this finding. Active learning could function as a barrier when participants were required to complete extensive textbook reading to prepare for class. Because there was only a single source of media, this practice violates the universal design for learning guideline of providing multiple means of representation. As a result of providing only a textbook, students who may struggle to read efficiently are disadvantaged when it comes to class preparation, which can negatively affect their experiences working with peers.

Universal design for learning endeavors to create learning experiences and spaces that are accessible to the greatest number of students possible. Employing universal design for learning may support the learning of all students in a course, not only students with disabilities. Furthermore, the use of universal design for learning aligns with the social model of disability. The social model of disability posits that individuals with impairments do not experience disability due to their biological differences, but instead due to the societal expectations placed on that individual. By employing universal design for learning, instructors are taking steps to create learning experiences that proactively address many potential barriers. Intriguingly, many of the ways in which active-learning practices were perceived to support perceptions of learning aligned with checkpoints and principles of universal design for learning. For example, Carson explained that group work helps him feel motivated to learn the material because he is no longer interested in earning straight A’s in his STEM courses. Group work for Carson seemed to align with the guiding principle of multiple means of engagement. In the future, exploring how universal design for learning and active learning relate to one another may help refine our understanding of these broad educational constructs.

### Self-advocacy in active-learning STEM courses

Universal design for learning can help instructors create more accessible educational experiences for students, yet using universal design for learning does not circumvent the need for individual accommodations within a classroom (Burgstahler, 2009). For this reason, it is important to understand the possible effects of active learning upon the self-advocacy of our participants. Not all participants considered active learning to influence their self-advocacy. Some participants perceived their need for self-advocacy to increase in STEM courses using active-learning practices primarily due to issues of overlooking accommodations and for the increased number of peer interactions during group work. Interestingly, although some participants acknowledged that they may have an increased need to engage in self-advocacy because of active-learning practices, they tend to feel more comfortable communicating with their instructor because they perceive the classroom to be more student-centered. The influence of active learning on self-advocacy and the examples of self-advocacy in active-learning STEM courses found in our study are summarized in Figure 4.

**Figure 4.**
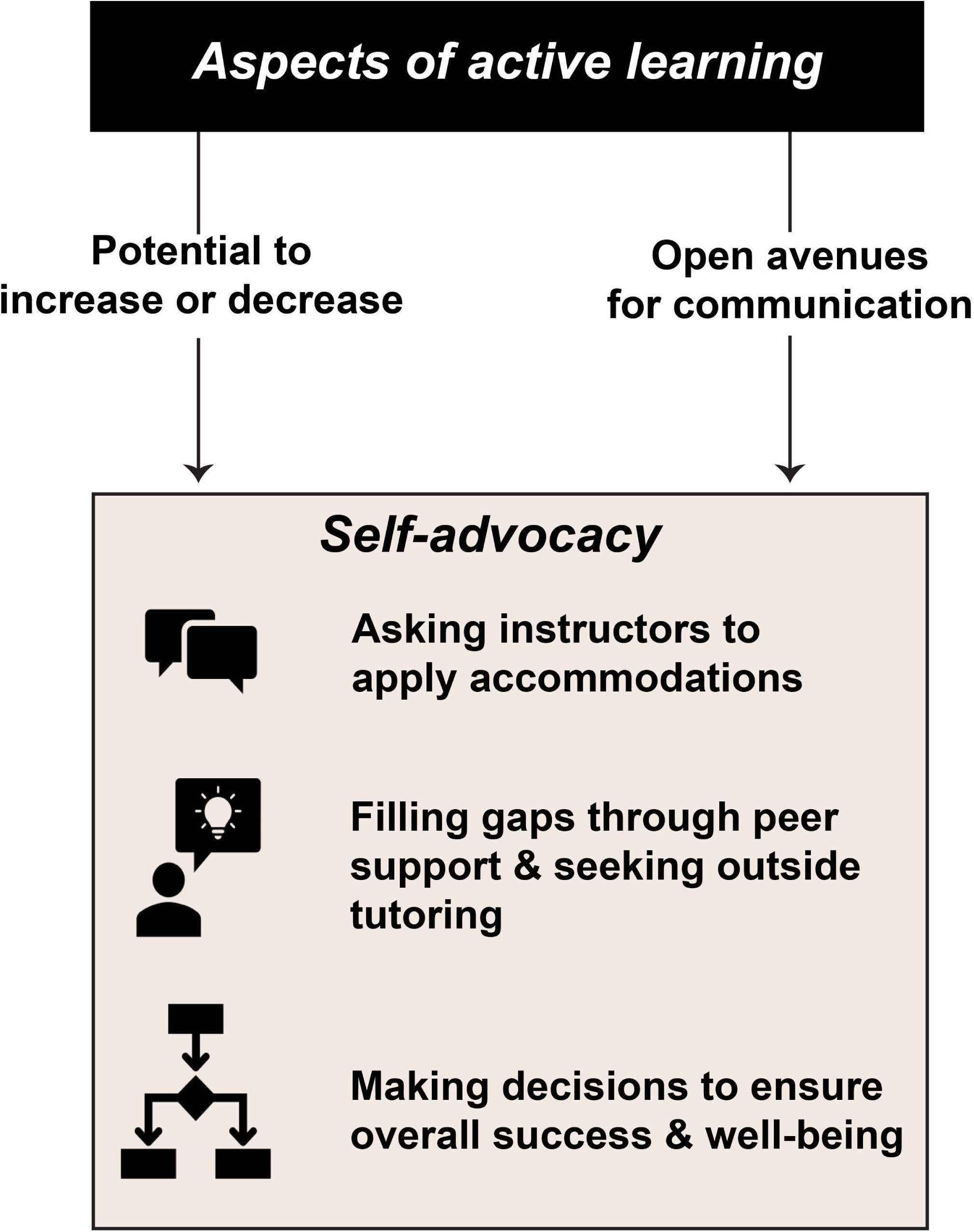
Summary describing how aspects of active learning influence self-advocacy for students with ADHD/SLD in our study. The tan box includes the examples of self-advocacy reported in our study: asking the instructor to apply accommodations, filling gaps through peer support and by seeking outside tutoring, and making decisions to ensure overall success and well-being.

### Implications for research

Disability is part of the human experience, but much existing higher education research ignores disability as a possible feature of the student experience (Peña, 2014). There is a need to consider the experiences of students with disabilities within future active-learning research. Our results are related to issues of fidelity of implementation. Fidelity of implementation is defined as “the extent to which the critical components of an intended educational program, curriculum, or instructional practice are present when that program, curriculum, or practice is enacted” (Stains and Vickrey, 2017, p. 2). The way in which instructors “in the wild” implement a particular instructional practice can differ from the implementation intended by the developers of the practice (Dancy et al., 2016; Offerdahl et al., 2018; Stains & Vickrey, 2017). In the context of STEM education, fidelity of implementation can be obstructed when a particular instructional strategy is enacted in a manner that neglects the critical components, or the essential elements, of that practice (Offerdahl et al., 2018). Yet defining the critical components of a particular active-learning practice is often not straightforward (Eddy et al., 2015). For example, many active-learning practices do not have a single developer, nor are the active-learning practices consistently defined within the existing literature (Waugh & Andrews, 2020). This may make it challenging for STEM instructors to readily identify how they should implement a particular active-learning practice and if that practice is appropriate given their own teaching contexts (Eddy et al., 2015; Waugh & Andrews, 2020). We recommend that as the field begins to theorize and conduct fidelity of implementation evaluations of existing evidence-based instructional practices that students with disabilities are included in these research efforts. We see opportunities to systematically characterize how instructional practices affect students with disabilities within the qualitative arm of fidelity of implementation evaluations (Stains & Vickrey, 2017). By consciously including students with disabilities in future research efforts and in fidelity of implementation evaluations, we may be better positioned to understand how instructional practices are affecting the experiences of students with disabilities in undergraduate STEM courses.

The findings of our study offer implications for future self-advocacy research. Our participants made decisions to ensure their overall well-being and academic success in a course. We hypothesize that these types of decisions involve knowledge of self, knowledge of STEM learning contexts, and the self-advocacy belief of agency (Pfeifer et al., 2020). It is also possible that students with different levels of self-advocacy may be perceiving active-learning STEM courses in distinct ways. Developing an instrument to measure self-advocacy for students with ADHD/SLD in undergraduate STEM courses will be an important next step in disentangling this potential relationship. One goal of this line of research would be to optimize active-learning STEM courses to best support individual self-advocacy.

### Implications for teaching

Broadly, our study offers evidence that implementation of a particular instructional practice can result in negative experiences for students with ADHD/SLD in active-learning STEM courses. As instructors, we have a responsibility to consider how students may be affected by the implementation of a particular active-learning practice. We encourage readers to consult existing evidence-based teaching guides offered by *CBE-LSE*, such as the guide on group work (Wilson et al., 2017), as well as other resources like the Practical Observation Rubric To Assess Active Learning (Eddy et al., 2015), and to share these resources with their colleagues to support more inclusive, and likely effective, implementation of active-learning practices. The language we use to talk about teaching matters. Taking time to clarify active-learning terminology and the essential elements of a teaching practice when discussing instruction may be one step towards promoting more effective implementation of that practice within our own departments (Dancy et al., 2016). Enhancing the implementation of active-learning practices could better support the learning of students with ADHD/SLD in undergraduate STEM courses.

Participants shared suggestions, including teaching recommendations, that they perceived would enhance their own experiences of learning in STEM courses using active-learning practices. We refer readers to Figure 3 (Supplemental File 6) to review these suggestions. We also discuss these suggestions in more detail below.

- **Consider student differences in your teaching.** Many participants explained that they find it frustrating when their instructors expect them to know something based on a single exposure to class content. When they ask their instructors questions, the instructor often states that they should already know the answer because the instructor explained it one time already during class. They also find it disheartening when instructors assume that they are lazy or have not put in the effort to reach a certain learning goal set forth by the instructor. Vivian’s quote in Figure 3 (Supplemental File 6) highlights that she wishes her instructors knew that she was putting in a lot of effort into her course work, but that sometimes her learning disability makes it challenging for her to meet the learning goals.
- **Know that how instruction is implemented directly affects participant success in a course.** Some participants in our study explained that they perceive themselves to be especially affected by how a STEM instructor teaches a STEM course. As discussed throughout our discussion section, how an active-learning practice is implemented matters. Therese spoke to this idea in her quote in Figure 3 (Supplemental File 6).
- **Explain your thinking to the entire class.** Our participants shared that hearing the instructor’s explanation is helpful for clicker questions, worked examples, and for more general directions to the class. Several participants shared that when the instructor does not explain their thinking or reasoning to the entire class, it hinders their learning. A lack of instructor explanation was a major barrier for our participant’s learning. Often when the instructor failed to reveal their own thinking to the class, participants would describe making decisions to seek third-party tutoring or to withdraw from the course. Kacey described why this type of situation is a negative influence on her perception of learning, in the context of clicker questions (Figure 3 and Supplemental File 6).
- **Provide interactive notes to support learning.** Our participants described how instructor-provided interactive notes were a major support for their learning. Some of our participants described that when an instructor is lecturing, the lecture can seem like disconnected thoughts or random words. When this happens, it can be difficult to identify key information to write in their notes. As a result, students may write irrelevant information in their notes that the instructor shares tangentially. This is problematic because the student may then use valuable time to study this irrelevant information. A few participants also shared that they may lose focus during class. When they regain focus, they are not aware of what information they have missed in the class. Providing interactive notes may help students to see where the class is now, and to identify what content they may need to follow-up on. Additionally, some of our participants had SLD in writing, which can make writing notes difficult. As we reported previously, nearly all participants in our study who qualified for a notetaking accommodation explained that they had difficulty using this accommodation (Pfeifer et al., 2020). Whether from not receiving notes at all, or receiving low-quality notes from the notetaker, many participants expressed that they would prefer to use their own notes. Providing interactive notes as a resource may allow students with ADHD/SLD to take more effective notes independently. Stewart explained that this type of resource would decrease his need for self-advocacy in his STEM courses (Figure 3 and Supplemental File 6).
- **Videos are preferred over extensive reading from the textbook.** One of the most profound barriers experienced by our participants was challenges in completing extensive readings required for flipped courses or in-class engagement activities. Our participants tended to speak more favorably for well-designed videos to learn course content. Bryce, a student who qualifies for alternative textbooks, discussed this more in his quote in Figure 3 and Supplemental File 6. Participants also appreciated videos that were well-aligned to the course because they could easily revisit the video to clarify course material at their discretion.
- **Add a roadmap for accommodations in the syllabus.** With implementation of active-learning practices comes questions of how accommodations will be administered within a STEM course (Gin et al., 2020). One of our participants, Erik, suggested that instructors provide directions for how accommodations are implemented within an active-learning STEM course that goes beyond the general disability statement often seen in syllabi (Figure 3 and Supplemental File 6). Most disability statements found in syllabi are the suggested statements from campus DRCs. These generic statements direct students who plan to request accommodations to contact the DRC and provide contact information for the DRC. Frequently, these statements do not provide more directions about the protocols the instructor uses to administer accommodations in the course. For instance, students must engage in additional communication (an example of a self-advocacy behavior) with their instructor to determine if the instructor prefers for them to take their exam at the DRC, or if the instructor plans to administer their extra time accommodations in the class. Students also must communicate with their instructor to learn how they can use their extended time accommodation for quizzes administered during class. Providing detailed accommodation practices with students in the syllabus may help students plan for accommodations. Alternatively, if instructors do not wish to include these protocols widely in the syllabus, having a prepared statement ready to share with students using accommodations could also serve a similar purpose.

We generated a comprehensive list of these recommendations from participants and our own recommendations (Table 5). Our researcher-generated recommendations were developed in response to the barriers and supports described by participants and as appropriate draw on existing suggestions from the literature. These recommendations are not intended to represent a panacea. Instead, they are offered so that instructors can work to address the needs of students with ADHD/SLD in active-learning STEM courses.

## Conclusion

We conducted foundational research to characterize how the implementation of active-learning practices in undergraduate STEM courses affects the perceived learning experiences of students with ADHD/SLD. Many participants expressed that they perceive themselves to learn best in an active-learning STEM course and explained how aspects of active learning support them. However, we also found many examples of how active-learning practices are perceived to hinder individual learning. Understanding these barriers can help STEM instructors become more aware of the potential pitfalls of active learning. Our results offer future directions to create more inclusive active-learning STEM courses. The development of more inclusive active-learning STEM courses is likely to support the retention of STEM students with ADHD/SLD, and other disabilities, within STEM majors.

## Supporting information

Supplemental Files Combined

## Acknowledgements

We thank our participants and members of the University of Georgia’s (UGA) Disability Resource Center (DRC) speakers’ bureau. We acknowledge Evie Reiter and McKenna Hendrickson for their contribution to preliminary analysis for this project. We are grateful to our present and past partners at the UGA DRC: Erin Benson, Sam Adair, and Pat Marshall. We also thank the Biology Education Research Group at UGA, especially Tessa Andrews, Peggy Brickman, and Paula Lemons, for their ongoing support and feedback through the lifetime of this project. We want to specially acknowledge Stephanie Halmo, Trevor Tuma, Emily Bremers, and Emmierose Scates for their helpful feedback on early iterations of this work.

This material is based upon work supported by the NSF Graduate Research Fellowship Program under Grant Number 1842396 (in support of M.A.P.). Any opinions, findings, and conclusions or recommendations expressed in this material are those of the author(s) and do not necessarily reflect the views of the NSF. This research was also supported in part by funds from the Division of Student Affairs at UGA (to J.D.S. and M.A.P.).

## Footnotes

1 We elected to use person-first language in our study, but we acknowledge that identity-first language is not the preferred terminology for everyone. Our participants tended to use person-first language when talking about themselves and person-first language is the preferred terminology for the American Psychological Association. For these reasons, we decided to use person-first language.

2 Learning disabilities in this study included dyslexia, ADHD, and autism.

3 SCALE-UP courses are implemented in classrooms that are uniquely designed to permit efficient group work with students typically sitting around circular tables containing computers or laptops with multiple projector screens and dry-erase boards arranged at the periphery of the room to facilitate sharing of information with the entire class (Beichner et al., 2007).

