## Supplemental Files Combined for "Promising or problematic? Perceptions of active learning from STEM students with ADHD and specific learning disabilities"

**Supplemental File Inventory**

| File Description | Page |
|---|---|
| **Supplemental File 1.** Screening survey questions. | 2 |
| **Supplemental File 2.** Interview questions. | 3 |
| **Supplemental File 3.** Coding matrix used for data analysis. | 4-5 |
| **Supplemental File 4.** Figure displaying active-learning aspects perceived to support or hinder participant learning. This figure depicts the same information as shown in Table 4. | 6 |
| **Supplemental File 5.** Table summarizing the influence of active-learning aspects on participant self-advocacy. This table displays the same information as shown in Figure 2. | 7 |
| **Supplemental File 6.** Participant suggestions for STEM instructors about active learning. This table displays the same information as shown in Figure 3. | 8-9 |

**Supplemental File 1.** Screening survey questions used for the study.

1. What is your major?

2. What year are you in school?

3. Are you 18 years of age or older?

4. What is your disability?

5. Have you taken a STEM course or are you currently enrolled in a STEM course for Fall 2018 that meets either the Science or Quantitative Reasoning requirement?

   Note: A list and a website link to a list of these courses were provided to participants.

6. Type in all the STEM courses you have taken or are currently enrolled in for Fall 2018.

7. In your most recent STEM course, did your instructor use active learning?

8. Select which examples of active learning you remember your instructor using in your most recent STEM course.

9. If you chose other in the question above, what other active learning did you do in your most recent STEM course?

**Active learning description provided in survey**

**Active learning** is a type of instruction that instructors in college science, technology, engineering, and mathematics (STEM) courses sometimes use. Active learning may occur when the instructor is not lecturing. *Examples of active learning are clicker questions, group work, completing worksheets in class either individually or in a group, class discussions, and student presentations*.

Note: Survey questions related to participant name and preferred contact methods are redacted.

**Supplemental File 2.** Interview questions used for the study.

Interview questions previously published are redacted. Bold font indicates interview questions yielding participant responses that were included in our structural coding process. Relevant data from other parts of our previously published interview protocol were also included in the structural coding process. That is, participant responses to these bolded interview questions were included as a minimum. Some participant responses included additional segments related to active learning, which emerged organically at other points in the interview. From these compiled segments (identified by structural coding) additional coding was done by the research team.

1. In your survey response, you mentioned that you have taken a STEM course to meet the Science and/or Quantitative Reasoning Core Curriculum requirement. **Which course(s) did you take?**

*To prepare for the interview, the interviewer compiles the list of courses the participant completed in Fall 2018. As the participant recalls these courses in the interview, the interviewer writes these courses on different notecards. The notecards are then used for Question #2.*

2. **Tell me about the type of instruction used in this course?**
    a. Did the instructor do a lot of lecture?
    b. Did you do any form of active-learning?

*Show the participant the cards from Question 1. Ask the student to describe the learning activities they remember for the course. Provide hand-out to participant with the various types of active-learning examples. Write active-learning practices on one half side and lecture activities on the other. After discussing all the notecards (courses), ask the participant to choose one STEM course they remember well that used mostly lecture, and one STEM course they remember well that used mostly active-learning practices. Use these courses as the basis for Questions 3 and 4.*

3. Think back to your STEM course that uses mostly lecture.
    a. Walk me through what a typical class was like for you.
    b. Tell me the specific ways you self-advocated in this class.
    c. Tell me about your interactions with your instructor.
    d. Tell me about your interactions with your peers.

4. **Think back to your STEM courses that used active learning…**
    **a. Walk me through what a typical class was like for you.**
    **b. Tell me the specific ways you self-advocated in this class.**
    c. Tell me about your interactions with your instructor.
    d. Tell me about your interactions with your peers.

5. **Do you learn better in a STEM course that uses lecture or active learning?**

6. **Do you think the type of instruction used in a STEM course influences your self-advocacy?**

7. **Is there anything you would like others (instructors, DRC coordinators, your peers,) to know about what it's like for you when you are in an active-learning STEM course?[1] If so, what?**

[1] The interviewer sometimes asked this question and omitted the phrase "active learning." For this reason, only participant responses related to active-learning STEM courses were considered for further analysis.

**Supplemental File 3.** Coding matrix used for data analysis.

**Date Coded Together**:

| **Matrix for holistic coding_Participant name** | |
|---|---|
| Question | Coder responses |
| What is the participant's preferred learning method in a STEM course? | |
| What supports their learning in an active-learning STEM course? | |
| What barriers do they encounter in an active-learning STEM course? | |
| Do they suggest a way to overcome the barrier? | |
| Did they withdraw from an active-learning STEM course? | |
| Does the type of course influence their self-advocacy? | |
| What do they want STEM instructors to know about their experiences in active learning courses?<br><br>Note: Make sure interviewer asked about active learning in the question. | |
| Top 2-5 quotes from this interview: | |

| | |
|---|---|
| Where would the participant fall in this positionality graphic? | 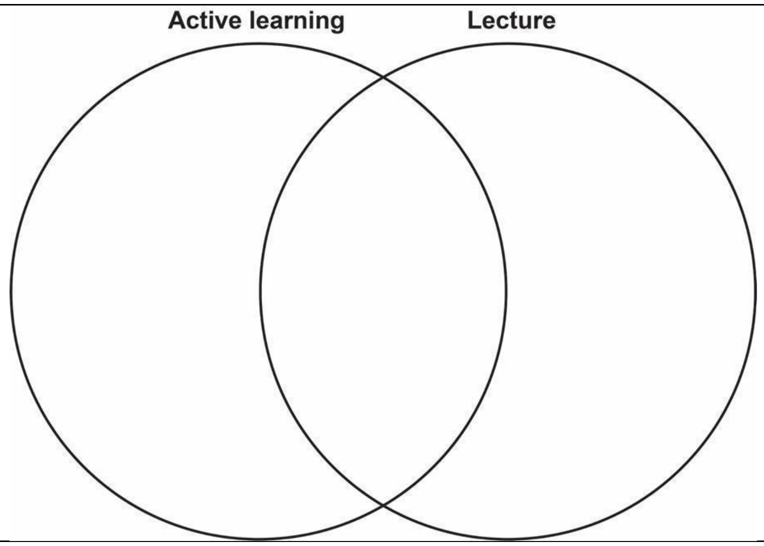  |
| Rationale for placement on graphic | |
| One-paragraph summary of participant’s perception of active learning | |

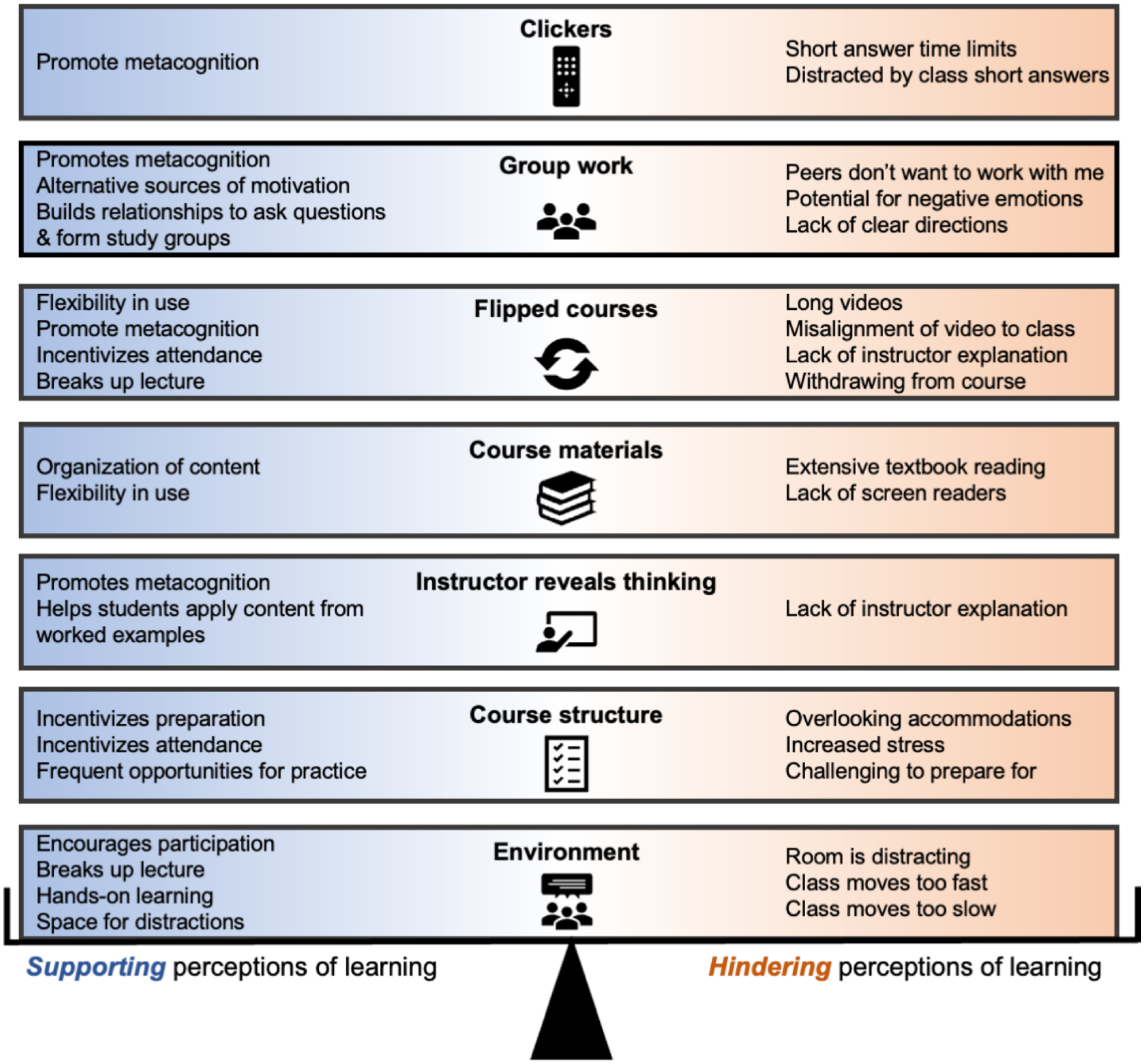


**Supplemental File 4.** Figure displaying active-learning aspects perceived to support or hinder participant learning. The aspects of active learning discussed by participants included: environment, course structure, instructor reveals thinking, course materials, flipped courses, group work, and clickers. Supports for flipped courses frequently overlap with other supports, a few of these key supports are indicated in the figure. Blue indicates aspects of active learning which were perceived to support the learning of participants. Orange represents aspects of active learning which were perceived to hinder the learning of participants. This figure depicts the same information as shown in Table 4.

**Supplemental File 5**. Summary of how aspects of active learning influence participant self-advocacy with example quotes. Table displays same information as shown in Figure 2.

| *Influence* | *Why?* | *Example quote* |
|---|---|---|
| **No influence on self-advocacy** | Participants see accommodations as always sufficient | If I was having trouble with a certain style of teaching, I probably wouldn't say anything because my accommodations are what's supposed to put me on an even playing field, right, so it's like I don't feel like I deserve any extra special stuff on top of it. —Sadie |
| **Decreased need for self-advocacy** | Participants consider active learning supportive | When I'm in an active learning kind of [course]… I understand the stuff more… I'm not usually put in a situation where I have to go up to the teacher and tell them, "I'm struggling really badly with this." Because we've already incorporated [active-learning practices] in there…I understand [the material] better. –Therese |
| **Increased need for self-advocacy** | Participants with unmet accommodation needs | I'm not getting extra time quizzes, and we have one before every class. It's a five-minute quiz. I would like two and a half extra minutes. Even reading, I read it, but I don't internalize it, so I gotta do it again, and then I gotta internalize it. I'll just email her. I emailed her recently about the extra time on the quizzes. She hasn't responded. —Kacey |
| **Increased need for self-advocacy** | Participants say group work is a situation that requires more self-advocacy than a lecture course | When you are more in a group situation you need a little bit more of if then just a basic lecture. In a lecture you can kind of come and go out and not have to do anything, not have to interact... I feel like when you're with others you want to talk more about yourself and lean more towards using more self-advocacy. Definitely in active learning situations you use more of it. —Brett |
| **No influence on self-advocacy** | Participants describe always practicing self-advocacy, no matter the course. | It [active learning] doesn't change the fact that I'm still being a self-advocate. —Erik |

**Supplemental File 6**. Participant suggestions for STEM instructors about active learning. Table displays same information as shown in Figure 3.

| ***Suggestion*** | ***Example quote*** |
|---|---|
| **Consider student differences in your teaching** | You have a certain bar set and if some students are meeting that, then you think that, that's an appropriate bar but if other students aren't, it's like are they not putting in enough effort? Or maybe they have a learning disability that's keeping them from getting up to that [goal] and you don't really know. —Vivian |
| **Know that how instruction is implemented directly affects student success** | My grade in a course is usually pretty highly affected by how the teacher teaches, and the quality of their teaching. So the better they teach, the better I'm going to understand it. And I think that active learning just works a lot better in general for me when implemented correctly. It really does make a huge difference. —Therese |
| **Explain your thinking to the entire class** | It's not right for when everyone gets the question wrong and it's very obvious, it's like 80% for answer D, and the answer was A. You just say worrisome things, and groan and moan, and [say] "Uh-oh, you all have a test on Wednesday, uh-oh." That's not helping anyone. That's not building up anyone. That's giving everyone anxiety. That's giving everyone a little more stress about not understanding something that we don't even know what [the instructor is] referring to exactly. There's so many parts of a question, we don't even know where people could potentially be going wrong…That doesn't give people hope or determination to figure a question out. –Kacey |
| **Provide interactive notes to support learning** | I would have less of reason to self-advocate if I had those resources. —Stewart |
| **Videos are preferred over extensive reading from the textbook** | [High-quality] videos work a lot better than trying to flood [read] a textbook, just passages and passages of [STEM content] in really, really big words. It works a lot better. –Bryce |

| | |
|---|---|
| **Add a roadmap for accommodations in the syllabus** | With regards to accommodations, they could always have a footnote about accommodations and just protocol. I think that would be helpful for students with disabilities. Because then they see okay, I have a roadmap. I can approach the professor and, it'd just be something that would be nice to have in a syllabus. –Erik |
